# Transient DREADD manipulation of the dorsal Dentate Gyrus in rats impairs disambiguation of similar place-outcome associations

**DOI:** 10.1101/2024.02.20.581173

**Authors:** Judith Lim, A. Souiki, P. Ahmad, Charlotte A. Oomen, Gerjan Huis in ’t Veld, Carien S. Lansink, Cyriel M.A. Pennartz, Umberto Olcese

## Abstract

The dentate gyrus subfield of the hippocampus is thought to be critically involved in the disambiguation of similar episodic experiences and places in a context-dependent manner. However, most empirical evidence has come from lesion and gene knock-out studies in rodents, in which the dentate gyrus function is permanently perturbed and compensation of affected functions via other areas within the memory circuit could take place. The acute and causal role of the dentate gyrus herein remains therefore elusive. The present study aimed to investigate the acute role of the dorsal dentate gyrus in disambiguation learning using reversible inhibitory DREADDs.

Rats were trained on a location discrimination task and learnt to discriminate between a rewarded and unrewarded location with either small (similar condition) or large (dissimilar condition) separation. Reward contingencies switched after a reversal rule, allowing us to track the temporal engagement of the dentate gyrus during the task. Bilateral but not unilateral DREADD modulation of the dentate gyrus impaired the initial acquisition learning of place-reward associations, but performance rapidly recovered to control levels within the same session. Modelling of the behavioural patterns revealed that reward learning and reward sensitivity were not associated with the DREADD-dependent impairment during acquisition learning, suggesting that either the ability to encode place-reward associations, or the fine-grained coding of place were instead affected. Our study thus provides novel evidence that the dorsal dentate gyrus is acutely and bilaterally engaged during the initial acquisition learning of ambiguous place-reward associations, although the exact neural mechanisms supporting this function still need to be fully understood.

## 1. Introduction

Remembering places associated with salient events and keeping these memories distinct when encountering similar spatial contexts are important cognitive functions to guide spatial navigation and decision making. The hippocampus is critically involved in memory (Eichenbaum et al., 1996), spatial navigation (O’Keefe, 1971; O’Keefe, 1976; O’Keefe and Nadel, 1978; McNaughton et al. 1996), and spatial memory (Eichenbaum et al. 1990; McDonald and White, 1995). Specifically, the hippocampus encodes individual places, trajectories and environments and their temporally associated sensory, motivational and more abstract events (e.g. place-reward associations) in a context-dependent manner (Leutgeb et al. 2007; Lansink et al. 2009; reviewed in Lisman et al. 2017; Latuske et al. 2018; van Dijk and Fenton, 2018). The mapping of highly similar contexts onto distinct neural representations by the hippocampus is crucial to successfully discriminate between contexts and is a process referred to as pattern separation. The ability to disambiguate similar experiences is thought to be degraded in the early stages of Alzheimer’s disease (Ally et al. 2013; Wesnes et al. 2014; Zhu et al. 2017; Leal and Yassa, 2018; Lee et al. 2020; Parizkova et al. 2020; Laczó et al. 2021), emphasizing the need to understand the neural basis of pattern separation.

The dentate gyrus subfield (DG) of the hippocampus is thought to be critically involved in pattern separation of similar contexts by orthogonalizing afferent input activity patterns into separate, distinct neural representations (Marr, 1971; Treves and Rolls, 1994; Hunsaker and Kesner, 2013; Knierim and Neuneubel, 2016). Empirical evidence for this disambiguation function attributed to the DG has mostly come from behavioural lesion studies in non-human primates (Hampton et al. 2004; Lavanex et al. 2006) and rodents (Gilbert et al. 2001; Hunsaker and Kesner, 2008; Morris et al. 2012; Lee and Solivan, 2009), as well as gene knock-out studies in rodents (McHugh et a. 2007; Kannangara et al. 2015; Yun et al. 2023). In these studies, dysfunction of the dorsal DG (dDG) was typically associated with an impaired ability to discriminate between adjacent positions (Gilbert et al. 2001; Hunsaker and Kesner, 2008; McHugh et a. 2007; Morris et al. 2012; Kannangara et al. 2015; Oomen et al. 2015; Yun et al. 2023) and object locations (Lee and Solivan, 2010), with animals making more mistakes in identifying the location associated with reward. Although the general conclusion from these studies is that dDG dysfunction impairs the ability to spatially separate the stimulus locations from one another, one could also argue that the deficit may, alternatively, lie in an impairment in forming the place-outcome association or the fine-grained encoding of place herein. To the extent of our knowledge, it thus remains unclear how the DG contributes to the acquisition of place- and outcome-related processes underlying spatial discrimination learning.

Specifically, it is unclear whether the DG contributes to actively maintaining recently acquired spatial representations (i.e. working memory), building up new spatial representations (e.g. spatial coding), acquiring place-outcome associations (i.e., conditioned place preference), or adjusting pre-established place-outcome associations (e.g. spatial reversal learning). Furthermore, the causal and immediate engagement of the DG in spatial discrimination remains elusive, because permanent damage to the DG in lesion and knockout studies may result in long-term compensation of DG function via other areas (Zelikowsky et al. 2013; Hong et al. 2018; reviewed in Vaidya et al. 2019).

To address these questions, we tested rats on a location discrimination task that we specifically developed to distinguish potential functions of the DG in spatial discrimination and representation, associative place-outcome learning and reversal learning abilities. To overcome the limitations of permanent lesion studies, we transiently disrupted the dDG with an inhibitory Designer Receptors Exclusively Activated by Designer Drugs compound (DREADDs; Roth et al. 2016) during the task. This targeted the excitatory cell population in the DG, consisting of granule cells and mossy cells, both densely projecting to area CA3. We found a significant impairment of discrimination performance of close (i.e. similar) but not distant (i.e. dissimilar) stimuli when the DG was disrupted, but no impairment of spatial reversal learning. Specifically, the DG-dependent discrimination deficit was present only during the acquisition phase of the task and recovered to control performance levels within a test session. Moreover, animals with an induced dysfunction of the DG were found to alternate more in their choice behaviour, independent of trial outcome. Our findings suggest that the DG is necessary during the acquisition of place-outcome associations to solve behavioural tasks that require fine-grained separation of similar spatial stimuli.

## 2. Methods

### 2.1. Animals

Twenty-three male Lister-Hooded rats (Envigo, the Netherlands) aged 7-8 weeks at arrival were housed in pairs in standard polycarbonate cages on a reversed 12-hour day-light cycle. Animals were habituated to their home cages with *ad libitum* water and food and were handled daily by the researchers for a period of 1 week. To motivate animals to perform the behavioural task, animals were food restricted to maximally 85% of the standard growth curve as obtained under *ad libitum* feeding conditions and had *ad libitum* access to water. All experimental procedures were performed in accordance with the European Directive 2010/63/EU, guidelines of the Federation of European Laboratory Animal Science Associations, the NIH Guide for the Care and Use of Laboratory Animals and were approved by the national and institutional ethics committees.

### 2.2. Behavioural task with DREADD manipulation

To probe spatial pattern separation, animals were trained and tested on a location discrimination (LD) task (Fig 1A). The LD task was adapted from the touchscreen LD operant platform task for testing hippocampus-dependent spatial memory in both rats (McTighe et al. 2009; Svensson et al. 2016) and mice (Clelland et al. 2009, Creer et al. 2010, Coba et al. 2012). In our LD task, the reward wells were positioned underneath each of the 8 stimulus loci presented on a screen, instead of a sole reward well located at the side opposite to the screen wall (Fig 1A; McTighe et al. 2009). This design reduced training times of rats (C.A.O., unpublished observations), likely because the spatially aligned stimulus and reward locations allowed for a more rapid formation of spatial stimulus-outcome associations. Moreover, rats performed self-initiated trials, facing the screen using a central nose poke well to ensure that animals were clearly exposed to the stimuli at the trial onset.

**Figure 1.**
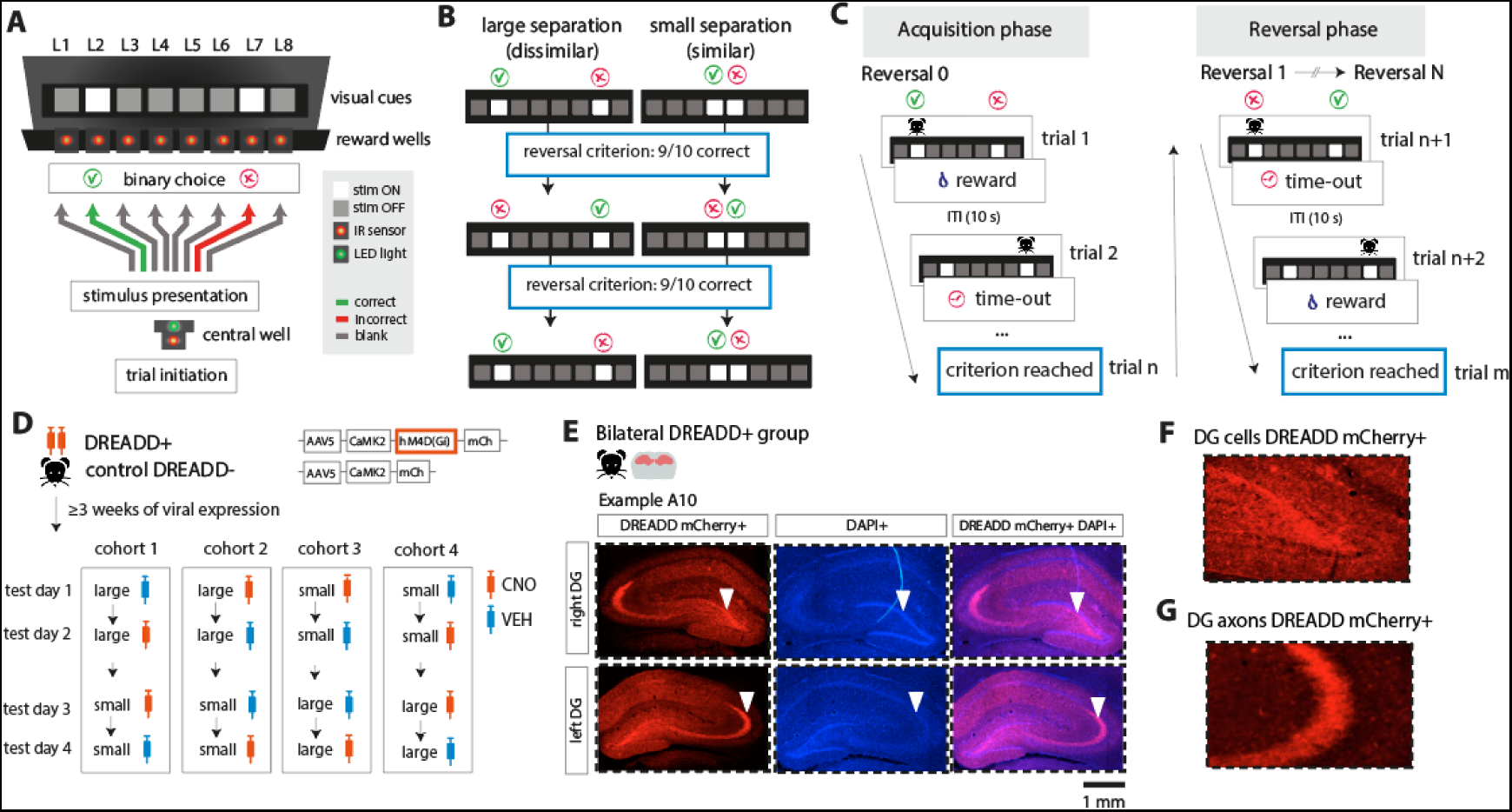
Task design of location discrimination task with chemogenetic silencing of dorsal dentate gyrus. **A)** Schematic overview of operant box with the CS+ and CS-visual cues (white squares) presented on a black monitor screen. After trial initiation at the central well, animals made a binary choice (left or right) with a nose poke in the corresponding cued nose poke well. **B)** Cued stimuli were presented with a large separation or small separation. After 9/10 consecutive correct choices, the reward contingencies switched. **C)** Schematic illustration of the learning phases of the task. At acquisition, animals identified the rewarded location by trial-and-error with positive (sucrose reward) and negative (time-out) feedback. In the reversal phase, animals re-learnt to identify the reversed target locations by adapting their choice behaviour. **D)** Chemogenetic targeting of the DG. Animals were bilaterally injected in the dDG with DREADD-mCherry (DREADD+ group, N=20) or control-mCherry (control DREADD-group, N=3). At testing, animals were injected with either CNO or saline (VEH) 30 min. before testing in a balanced within-subject design. **E)** Histological verification of DREADD-mCherry+ in dDG in bilateral DREADD+ group (N=11). Example animal (A10) with bilateral DREADD expression in dDG (left panel), DAPI expression (middle panel) and overlaid (right panel). White triangles indicate subregions of the hippocampus expressing DREADDs. **F)** Close up of DREADD-mCherry+ expression in DG cells and **G)** DG-projecting axons in CA3.

Animals were trained in an automated operant box (52 cm width × 40 cm height × 50 cm depth) with three black aluminium walls attached to a black monitor screen (64 cm width × 3 cm depth, Liyama Prolite B2776HDS) and a metal bar floor with 1 cm spacings. A black polyoxymethylene strip with 8 reward wells (each 5 cm in height × 6 cm wide × 4 cm depth) was located below the screen, so that 0.1 ml 15% sucrose water rewards could be delivered from syringe pumps via plastic tubes. Identical visual cues were presented on the screen as white squares (2 cm width × 2 cm height, center-to-center stimulus distance = 6 cm) at 8 possible stimulus locations corresponding to the reward wells located below the screen. Stimuli were generated with Psychtoolbox-3 in MATLAB (Brainaird, 1997) and the task setup was controlled with custom-written scripts in MATLAB.

#### 2.2.1. Behavioural pretraining

In the initial LD pretraining phase, animals learnt to respond selectively and reliably to 1 out of 8 possible cued locations (L1-L8, see Fig 1A) and the rewarded location was randomly chosen out of those 8 sites every trial. At trial start, a green LED light turned on at the central well, after which the animal could initiate the trial with a nose poke into the central reward well for at least 0.3 sec. After successful trial initiation, animals received a sucrose water reward with 20% probability to motivate the animal to self-initiate trials. If the animal poked correctly at the cued location (“correct trial”), a sucrose water reward was given at a reward well located below the corresponding cue stimulus. However, if the animal poked incorrectly, viz. at an uncued nose poke location (“incorrect trial”), a white LED light placed at the wall opposite to the nose poke panel was lit for 3 sec followed by a 5 s time-out. The visual cue disappeared after the choice was registered by a nose poke sensor located at each corresponding reward well. After the choice period, the trial ended and an inter-trial-interval (ITI) of 10 sec followed, during which the monitor screen was kept black to maximize the saliency of the white cue stimulus during trials. To proceed to the next phase of the experiment, the animal had to finish 90 trials within one session of 40 min with a performance of minimally 80% correct trials for two consecutive sessions.

#### 2.2.2. Viral injection surgery

After completing the pretraining, one batch of animals (N=21 rats, DREADD+ group) was bilaterally injected in the dDG with pAAV5-CaMKIIa-hM4D(Gi)-mCherry (virus titer ≥ 3×10^12^ vg/ml, Addgene, North Carolina, USA). Inhibitory DREADDs are activated by the artificial ligand clozapine-N-oxide (CNO; Hsiang et al. 2014; Roth 2016; Mathos et al. 2019; Visser et al. 2020; Domi et al. 2021; Lesuis et al. 2021; Visser et al. 2022; but see Gomez et al. 2017) and transiently manipulate neural activity in targeted neurons for about two hours following CNO injection (Roth, 2016). To control for confounding CNO effects on motor activity and cognition due to CNO being converted into clozapine (i.e., unrelated to DREADD receptor activation; McLaren et al. 2016; Gomez et al. 2017; Manvich et al. 2018), a separate batch of animals (N=3 rats, control DREADD-group) were bilaterally injected in the dDG with pAAV5-CaMKIIa-mCherry with no DREADD compound (virus titer ≥6×10^12^ vg/ml, Viral Vector Facility, ETH Zürich, Switzerland). The surgical procedures were identical for the DREADD and control groups and were performed in line with standard operating procedures for intracerebral viral injections. Animals received a subcutaneous injection of buprenorphine (0.01-0.05 mg/kg) and meloxicam (2 mg/kg) 20-30 min prior to anaesthesia. Animals were anaesthetized with isoflurane (induction level: 3-5% isoflurane; maintenance level: 0.5-3% isoflurane) and injected with 0.48 μl of either the DREADD or control viral vector per injection site at a flow rate of 0.056 μl per min and 10 min post-injection wait-time. Injections were made with an automated nanoliter injector (Nanoject II, Drummond Scientific Company, USA) at the following anterior-posterior (AP), medial-lateral (ML) and dorsal-ventral (VL) coordinates relative from Bregma: -2.7 mm AP, ±1.2 mm ML, -3.7 mm DV in the anterior dorsal DG; -3.7 mm AP, ±2.0 mm ML, -3.1 mm DV in the posterior dorsal DG. For the spread of the virus in the DG, see section 3 (Results). After surgery, animals were solitarily housed for 3-7 days with soft food and water *ad libitum* provided.

#### 2.2.3. Behavioural training

After recovery from viral injection surgery, animals were housed in pairs again and re-trained on the pretraining paradigm until they reached the level of performance observed before surgery. Animals were then advanced to the training phase, in which they learnt to discriminate between a rewarded and an unrewarded stimulus location with medium separation distance (18 cm). Animals initiated each trial similarly as in the pretraining phase, whereafter animals were presented with two visual cues and learnt to identify which of the two locations was rewarded by means of trial-and-error: the animal received a sucrose water reward if the choice was correct and a time-out of 5 s if the choice was incorrect, cued by LED illumination of the training box (as described for the pretraining phase). After performing 9 out of 10 consecutive correct trials, the reward contingencies reversed to assess reversal learning and avoid a spatial bias to one side of the box (Fig 1B-C). Animals were trained for 6 sessions, during which the locations of each unique stimulus pair was varied over sessions with a consistent separation distance (18 cm per location pair, medium separation): location 1 and 3 (L1– L3), 2 and 4 (L2–L4), 3 and 5 (L3–L5), 4 and 6 (L4–L6), 5 and 7 (L5–L7), and 6 and 8 (L6–L8).

#### 2.2.4. Behavioural testing with and without DREADD intervention

After completing training with medium distance, animals were ready for testing on the LD task with either small separation (L4-L5, distance = 6 cm) or large separation distance (L2-L7, distance = 30 cm) during neurointervention. Thirty to forty minutes before testing, animals received an i.e. injection with either the DREADD ligand clozapine-N-oxide (CNO; 3 mg/1ml/kg in saline, HelloBio.com) or saline (vehicle, VEH; 1 ml/kg) as a control treatment. Animals were tested on each combination of task separation and neurointervention in a total of four sessions (one session per test day, Fig 1D): small separation with vehicle (henceforth abbreviated as Small-VEH) or CNO (Small-CNO) and large separation with vehicle (Large-VEH) or CNO (Large-CNO). To counterbalance the impact of task manipulations on LD performance, animals were divided into four cohorts that were each tested on a different task manipulation on each testing day (Fig 1D). Procedures for the LD test sessions were the same as described for the LD training sessions, with the only difference that the maximum number of reversals was set to 6 within 60 min, so that animals had a similar testing exposure across sessions.

### 2.3. Histology

After experiments had been completed, animals received a lethal dose of 1 ml 20% euthasol via an i.p. injection and were transcardially perfused with 4% paraformaldehyde (PFA) in phosphate buffered saline (PBS) to verify virus expression in the dorsal DG. Brains were extracted and stored in a 4% PFA solution at 4°C for 2 days. To prepare for slicing, brains were immersed in 15% sucrose in PBS for 1 day and 30% sucrose in PBS (pH 7.3) for 2 days. Brains were sliced in 40 μm coronal sections using a microtome and stored overnight in PBS at 4°C. To stain fluorescent cells tagged with mCherry, sections were stained with DAPI (300 nM in PBS, Sigma). Sections were immersed in DAPI for 5 min at room temperature, after which DAPI was removed and washed 3 times with PB (pH 7.3). Stained sections were mounted on microscope slides (SuperFrost Plus, Thermo Scientific) and stored overnight at room temperature. Sections were imaged using a fluorescence microscope (Leica Microsystems DM300) at 10x magnification to capture viral expression in the dDG (Fig 1E-G, Suppl. Fig 1).

### 2.4. Data analysis

#### 2.4.1. Location discrimination and reversal performance

We first determined how task behaviour and locomotion activity were modulated by stimulus distance and DREADD manipulation of the DG during the LD task.

Location discrimination performance was quantified by the following behavioural measures. The number of trials that the animals needed to reach the reversal criterion of 9 out 10 consecutive correct choices was defined as the trials to criterion (TTC; Livesay and Bayliss, 1975; Oomen et al., 2013; McTighe et al. 2009), where low and high values of TTC indicate fast and slow learning of the reward contingencies, respectively. The proportion of errors was computed as the number of incorrect choices divided by the number of total choices and indicates how well the animal discriminated between the target and non-target cued locations. The response latency (in seconds) was defined as the time from stimulus onset until the animal poked at the target or non-target location (L4 and L5 for Small sessions; L2 and L7 for Large sessions; Fig 1B). The number of blank pokes was defined as the number of nose pokes that were made into wells that were never cued nor rewarded during the test sessions (L1, L3, L6 and L8 for both Small and Large sessions; Fig 1B). The number of blank pokes (i.e., at non-cued wells) was determined to probe how spatially precise animals poked at the target and non-target wells.

Each task performance measure was compared between the task separation and treatment conditions for each learning phase and virus expression group (bilateral DREADD+, unilateral DREADD+, control DREADD-). Measures were separately computed for each learning phase to contrast task performance across different learning phases, viz. the acquisition phase (Acq, where the animal did not adjust its behaviour to a first reversal yet) and the subsequent reversal phase, where the temporal rank order of reversals is indicated by R1, R2, …, R_n_. We fitted a generalized linear mixed model (GLMM) to each behavioural measure to account for the crossed random effects of the data (Equation 1; function *fit_glm*, MATLAB), since the model factors treatment and separation were crossed within each animal. Each GLMM was fitted with a set of distributions, including the normal, inverse gaussian, poisson, and gamma distribution, and the best model fit was determined as the model with the lowest sum of squared errors (SSE). We then performed an ANOVA on the model coefficients to test for the main and interaction effects of treatment and separation. Post hoc comparisons between the treatment conditions for each separation were made with a Wald test on the estimated marginal means (function *emmeans, MATLAB,* source code: github.com/jackatta/estimated-marginal-means), computed as the means of each model factor adjusted for the means of the other model factors. All post hoc p-values were adjusted with a false discovery rate correction for multiple comparisons. The significance level was set at p<0.05. Note that, in case of unbalanced group sized, estimated marginal means – that are however not biased by such imbalances – might not align with the observed means that we used in all plots.

The GLMM was specified in the general form as follows:

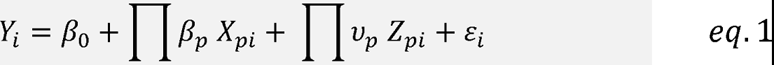

Here, *Y_i_* is the behavioural task measure corresponding to the levels of the predictor variables *X*_1*i*_, *X*_2*i*_, …, *X_pi_* and random effects variables *Z*_1*i*_, *Z*_2*i*_, …, *Z_pi_* of each trial *i*. The beta coefficients *β*_0_, *β*_1_, …, *β_p_* correspond to the predictor variables *X_pi_*, where *β*_0_ is the intercept. The random effects coefficients *v_p_* correspond to the random effect variables *Z_pi_*. Lastly, *ε_i_* indicates the residual error of the model fit for each trial *i*. The product terms of the predictor and random effects variables indicate the interaction term between the task variables separation x treatment. The weights of the coefficients are not described in the current study, as the coefficients differed for every model fitted to each DREADD expression group, task learning phase and behavioural task measure. As an example, the sample distribution of TTCs for the bilateral DREADD+ animals best fitted with an Inverse Gaussian distribution, where post hoc comparisons revealed a significant higher TTC for Small compared to Large sessions during the acquisition phase (*p*=0.043, Wald test; Fig 2F). Similar post hoc results were obtained with different distributions that fitted worse, e.g. fitting the data with a Gamma and Normal distribution yielded similar post hoc results (*p*=0.013 and *p*=0.011, respectively, Wald test).

**Figure 2.**
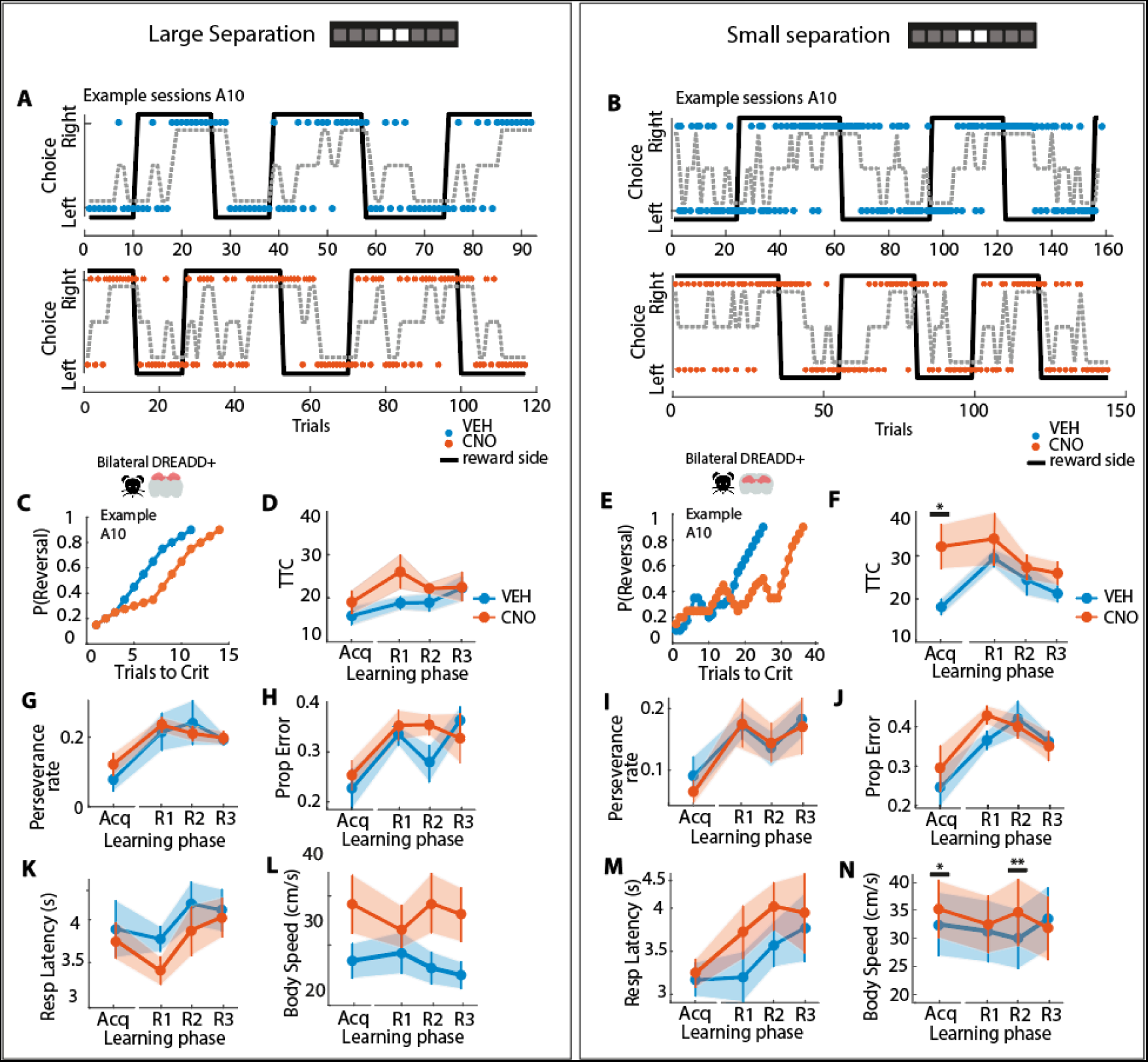
Task behaviour during location discrimination task for CNO and control treatment conditions. **A)** Example choice behaviour for a representative bilateral DREADD+ expressing animal (A10) with saline treatment (blue dots, top panel) and with CNO treatment (orange dots, bottom panel) for a Large separation session. Grey dashed lines indicate the smoothened choice behaviour (bin size = 4 trials). Black lines indicate the rewarded side (left or right). **B)** Same as A, but for a Small separation session. In all subsequent panels, left-side panels represent Large sessions (panels A, C, D, G, H, K, L) and right-side panel represent Small sessions (panels B, E, F, I, J, M, N). **C,E)** Reversal probability to make 9 out of 10 consecutive correct choices for the same animal (A10) as panels (A,B) with saline (VEH, blue line) and CNO treatment (orange line) across trials for the initial acquisition phase. **D,F)** Number of trials needed to reach the reversal criterion for the bilateral DREADD+ group (N=11) with VEH (blue line) and CNO (orange line) treatment for each learning phase. For (F), task performance was significantly impaired during the initial acquisition phase when DG was inactivated (GLMM ANOVA with post hoc Wald tests, p=0.043). **(G,I)** Proportion of trials where the same choice was made as the previous trial (perseverance rate) for each learning phase for the bilateral DREADD+ group. **(H,J)** Proportion of incorrect trials for each learning phase for the bilateral DREADD+ group. **(K,M)** Response latency (in seconds) for each learning phase for the bilateral DREADD+ group. **(L,N)** Body speed (in cm/s) during the choice epoch for each learning phase for the bilateral DREADD+ group. Error bars indicate average task variables pooled across animals (mean± SEM). Significance is indicated by * p<0.05, ** p<0.01, *** p<0.001.

#### 2.5.2. Locomotion activity during task behaviour

To control for non-specific DREADD effects on locomotion activity, we also quantified the rat’s body position from video data collected at 25 fps for animals with bilateral DREADD expression (N=7). Video frames were processed with DeepLabCut (Mathis et al. 2018), a semi-automated tracking program that uses deep neural networks to estimate the position of body parts based on manually labelled test frames. We labelled 6 body parts every 80-120 frames per animal, including the body center, left shoulder, right shoulder, neck base and snout. To obtain a reliable estimate of body position, we averaged the body part coordinates to a center-of-mass (X,Y) coordinate for every frame. Noise artefacts were reduced by removing those estimated coordinates that fell outside of the LD box, whereafter the tracking data was smoothed with a moving average filter (width: 10 frame samples) (function *smoothdata*, MATLAB) and an interpolation filter applied to a window of 100 msec (function *interp1*, MATLAB). Locomotion activity was quantified as body speed, defined as the travelled distance per second (cm/s) from the trial start until the trial end. The travelled distance was computed as the square root of the sum of squared differences between each pair of X coordinates and squared differences between each pair of Y coordinates. Body speed was compared between the four different combinations of separation and treatment conditions with a GLMM, separately for each reversal. Post hoc comparisons between treatment conditions were made with estimated marginal means.

#### 2.5.3. Fitting choice data to reinforcement learning models

To understand which aspects of spatial learning were affected by DREADDs, we estimated the learning parameters underlying place-outcome learning driven by reward using a validated reinforcement learning model (henceforth referred to as ‘RL model’; Rescorla & Wagner, 1972; O’Reilly and den Ouden, 2015; Sutton and Barto, 2018; Metha et al. 2020). Specifically, this model allowed us to investigate whether the learning of place-outcome associations was driven by reward, viz. by assessing the learning rate (*alpha* parameter) and 2) the tendency of animals to choose the rewarded location based on the reward value associated with each place (*beta* parameter). In turn, the quantification of these reward-related parameters may explain why we observed a selective dDG-driven deficit during the acquisition phase of the task (Fig 2D,F), when animals presumably acquired the place-outcome associations. Moreover, to determine whether animals persisted in their choice despite receiving reward in the previous trial, we extended the RL model with an additional perseverance parameter (*delta* parameter; henceforth referred to as Perseverance RL model, ‘PRL model’; adjusted from Metha et al. 2020).

To model how the choice behaviour of animals during the LD task was modulated by stimulus distance and DREADD manipulation of the DG, we fitted the choice data of each animal to the RL and PRL models. The models were then compared using the Akaike Information Criterion (AIC; Akaike, 1998) to determine which best fit the behaviour of each animal.

We first fitted the RL model to the choice data (Rescorla & Wagner, 1972; Metha et al. 2020; Sutton and Barto, 2018). This model was structured as follows. For each trial *t*, the probability of choosing the right stimulus (*P_right,t_*) was computed with a sigmoid function (Equation 2a) that models how the learned stimulus values (*V_right,t_* and *V_left,t_*) translated into a choice (Equation 2b). The sigmoid equation generally follows the behaviour observed in humans and rodents performing reversal tasks (O’Reilly and den Ouden, 2015) where the subject is more likely to choose the option with the higher stimulus value, but can also switch to the less valued choice from time to time by way of spontaneous alternation. The choice probability thus remains stochastic, i.e. without reaching values of 0 or 1. The value of each stimulus (*V_right,t_* and *V_left,t_*) was modelled with the Rescorla-Wagner equation (1972) that follows prediction-error based learning, in which the stimulus gains reward value when the outcome on the previous trial (*r*_*t*-1_) exceeds the predicted reward value on the previous trial (*V*_*t*-1_). Importantly, the prediction error scales with the learning rate *alpha*, that represents how quickly animals acquire the reward value when the stimulus value changes across trials (e.g. a low *alpha* indicates slow learning). We then estimated the slope parameter *beta*, that indicates how strongly the (binary) choice of the animal follows the stimulus value (graded) across trials (e.g. a low *beta* indicates that the animal chooses closer to chance level, rather than following stimulus value). For the Perseverence RL (PRL) model, the RL model was extended with a *delta* parameter that estimates the degree with which the animals persisted in the choice of the previous trial (Equation 2c). A positive value of *delta* indicates that an animal was more likely to respond on the same side as the previous trial (perseverance). Conversely, a negative value indicates that animals were more likely to switch from side to side on consecutive trials (alternation). In this model (Equation 2c) *C_left_* and *C_right_* are indicator variables, taking the value of 1 if the corresponding stimulus is chosen, and 0 otherwise. The parameters *alpha, beta* and *delta* were estimated for each animal based on the maximum likelihood fit to the choice data. We then compared the *alpha*, *beta* and *delta* parameters between treatment conditions for every separation condition with a GLMM as specified in section 2.4.1. All models were fitted with a constant offset value of 0.5 to reflect the choice probability at chance level at the start of each session. *P*-values were adjusted with an FDR correction for multiple comparisons. Significance levels were set at *p*<0.05.

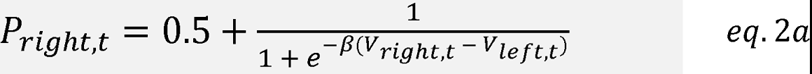

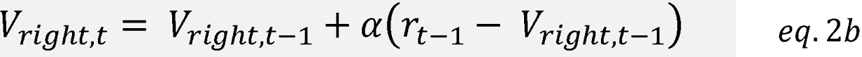

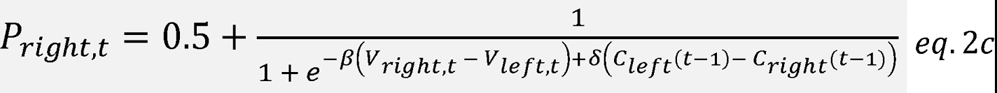

To study the role of the dorsal dentate gyrus (dDG) in disambiguating similar spatial representations, we selectively modulated the dDG with an inhibitory DREADD (AAV5-hM4D(Gi)-mCherry) in Lister Hooded rats (N=20) that performed a location discrimination (LD) task (Oomen et al. 2013). In this task, animals discriminated between a rewarded and an unrewarded location as indicated by a CS+ and a CS-visual cue (Fig 1A), where the separation distance was either small (i.e. similar location) or large (i.e. dissimilar location). Within each session, reward contingencies were switched after 9 out of 10 correct consecutive choices (‘reversal criterion’; Fig 1B-C) to avoid spatial biases for either cued location and to allow us to probe the spatial discrimination ability and cognitive flexibility required for adapting to changed reward contingencies. Animals were injected with either saline (vehicle treatment, VEH) or CNO (test treatment) and tested in each separation condition in a counterbalanced manner. To account for possible confounding effects on learning and motor behaviour due to CNO being converted into clozapine (i.e., unrelated to DREADD receptor activation; McLaren et al. 2016; Gomez et al. 2017; Manvich et al. 2018), we also tested a separate group of animals injected with a control viral vector lacking receptors for DREADDs (N=3).

Upon histological inspection, we separated animals in three groups based on the viral expression patterns in the dDG. We observed animals with DREADD-mCherry expression in both the left and right DG (bilateral DREADD+ group, N=11, Fig 1E-G) and animals with DREADD-mCherry expression in either the left or right DG (unilateral DREADD+ group, N=6, Suppl. Fig 1B). Although unilateral expression was not expected since the DREADD was injected bilaterally, we included this separate group of animals for further analysis to determine whether the discrimination ability of the dDG was lateralized in our task. Indeed, previous studies suggested that there may be a lateralization of spatial discrimination abilities in humans (Miller et al. 2018) and rodents (Song et al. 2020) due to perturbed communication between the left and right hippocampus (Zaidel 1995). The last group consisted of animals injected with the control virus lacking DREADD, showing bilateral mCherry expression in the DG as intended (control DREADD-group, N=3, Suppl. Fig 1C). In bilateral and unilateral DREADD+ animals, mCherry was clearly expressed in DG cell bodies: the granule cell layer and the hilus subregion were densely targeted, while dense axonal innervation was observed projecting from the DG to CA3 (Fig 1G; Suppl. Fig 1A-B). mCherry expression was most prominent in the dorsal part of the DG, extending from -1.72 mm to -4.56 mm AP from Bregma. mCherry expression was also observed in the CA1 layer bilaterally in 3 out of 20 rats (15%, Suppl. Fig 1A) and unilaterally in 5 out of 20 rats (25%, Suppl. Fig 1B), likely because the CA1 layer had to be penetrated to have the injections reach into the DG. mCherry expression was also observed bilaterally in the subiculum layer in 3 out of 20 rats (15%) and unilaterally in 6 out of 20 rats (30%, Suppl. Fig 1A). A highly similar expression pattern was observed in the DG for the control animals lacking DREADD, showing dense mCherry expression in the granule cell layer and the hilus subregion (Suppl. Fig 1C). In this control group, mCherry expression was also observed in the subiculum layer unilaterally in 1 out of 3 rats (30%). We observed qualitatively similar task behaviour in rats co-expressing DREADDs in the CA1 and subiculum either unilaterally or bilaterally (data not shown) and therefore included these rats for further analysis. Together, DREADD-mCherry was mainly expressed in the dorsal part of the DG, but a co-involvement of areas CA1 and subiculum cannot be excluded.

### 3.1. Transient DREADD manipulation of dorsal DG cells impairs spatial discrimination learning but spares reversal learning

To discern changes in spatial discrimination and reversal learning, we quantified the task performance separately for the acquisition phase (Fig 1C, left panel) and subsequent reversal phase (Fig 1C, right panel). Task performance was probed as the number of trials needed to reach the reversal criterion (trials to criterion, TTC; Livesay and Bayliss, 1975; Clelland et al. 2009; McTighe et al. 2009; Oomen et al. 2013). On average, animals made 5.50 reversals (± 0.19 SEM) for Large sessions (Suppl. Fig 2A) and 4.78 reversals (± 0.36 SEM) for Small sessions (Suppl. Fig 2B). There was a main effect of separation on the number of reversals made by bilateral DREADD+ (*F*(40)=39.61, *p*=1.82E-07, GLMM; Suppl. Fig 2A) and control DREADD-animals (*F*(8)=8.16, *p*=0.021, GLMM; Suppl. Fig 2C), but not for unilateral DREADD-animals (*F*(20)=0.10, *p*=0.75, GLMM; Suppl. Fig 2B). This showed that animals tended to make less reversals when discrimination was more difficult. No effect of treatment was observed on the number of reversals for any group (bilateral DREADD+: *F*(40)=1.10, *p*=0.30, GLMM; unilateral DREADD+: *F*(20)=0.10; *p*=0.75, GLMM; control DREADD-: *F*(8)=1.09, *p*=0.33, GLMM), showing that animals remained able to perform the reversal task rule despite of chemogenetic manipulations.

To determine whether task performance depended on the separation distance, we first compared TTC between Small and Large sessions for each task phase for vehicle sessions only (see Suppl. Table 1). There was a main effect of separation on task performance for bilateral DREADD+ animals after the first reversal was made (*F*(19)=11.60, *p*=0.0030, GLMM; Suppl. Fig 2D), showing that animals performed worse in Small compared to Large sessions. No effect of separation distance was observed for the acquisition phase and subsequent reversals within the same session (*p*>0.05, GLMM; Suppl. Fig 2D). These findings suggest that animals performed worse after the first reversal, but rapidly improved their performance and reached similar performance levels within the same session regardless of discrimination difficulty. In contrast, there was no main effect of separation distance on task performance for unilateral DREADD+ (Suppl. Fig 2E) and control DREADD-animals across learning phases (*p*>0.05, GLMM; Suppl. Fig 2F). Together, these results show that the baseline performance of animals generally did not depend on the task difficulty, except for the bilateral DREADD+ group, which, in the absence of CNO injections, performed worse after the place-reward contingencies of similar loci reversed for the first time.

We next determined how the task performance was modulated by the treatment depending on the separation distance, where the TTC was compared between treatment sessions for each separation level and task phase separately (Fig. 2A-B; see Suppl. Table 1). For bilateral DREADD+ animals, there was a main effect of separation (*F*(40)=5.49, *p*=0.024, GLMM) and treatment (*F*(40)=5.75, *p*=0.021, GLMM) on TTC, where task performance significantly worsened during the acquisition phase for CNO compared to VEH treatment for Small sessions (*W*=2.40, *p*=0.043, Wald test; Fig 2E-F). No effect of treatment was observed for the subsequent reversal phase (*p*>0.05, GLMM; Fig 2E-F; Suppl. Fig 2M). In contrast, task performance was not different between treatment conditions for Large sessions at any task phase (*p*>0.05, GLMM; Fig 2C-D; Suppl. Fig 2J), suggesting that animals only had difficulty disambiguating stimuli when the locations were close in space. Unilateral DREADD+ animals showed no effect of treatment, regardless of the separation distance, at any task phase (*p*>0.05, GLMM and Wald test; Suppl. Fig 2K,N), suggesting that the unilateral targeting of the dDG did not impair spatial discrimination learning. Surprisingly, there was a main effect of treatment on TTC (*F*(7)=13.91, *p*=0.0074, GLMM) after the second reversal was made by control DREADD-animals, where task performance was lower following CNO injection compared to VEH for Small sessions (*W*=3.73, *p*=0.015, Wald test; Suppl. Fig 2L,O). Given the large spread in TTC across control animals for Small sessions however (Suppl. Fig 2O), we believe that this finding should be taken into account with caution. Together, our findings suggest that bilateral but not unilateral dDG dysfunction impairs the spatial discrimination ability during the acquisition of associations between similar locations and reward.

To determine whether other measures of task performance were affected by modulating the dDG, we also compared the proportion of incorrect choices (error proportion, inverse of proportion correct; Gilbert et al. 2001; Lee and Solivan, 2009; Oomen et al. 2013, 2015), the time needed to report the choice (response latency; Oomen et al. 2013), the tendency of the animal to persevere its previous choice (perseverance rate) and the spatial accuracy of reported choices (see Suppl. Table 1). For bilateral DREADD+ animals, there was no effect of treatment on these performance measures for each separation level and task phase separately (*p*>0.06, GLMM and Wald test; Suppl. Fig 3). For unilateral DREADD+ animals, there was a main effect of separation and treatment on response latencies after the fourth reversal (separation: *F*(9)=8.09, *p*=0.019; treatment: *F*(9)=9.11, *p*=0.015, GLMM) and fifth reversal (separation: *F*(5)=71.50, *p*=0.00038; treatment: *F*(5)=35.98, *p*=0.0018, GLMM). Specifically, response latencies were lower following CNO injection compared to VEH after the fourth reversal for Large (*W*=2.36, *p*=0.043; Suppl. Fig 3K) and Small sessions (*W*=3.02, *p*=0.029, Wald test) and fifth reversal for Small sessions (W=6.00, *p*=0.0037, Wald test; Suppl. Fig. 3L). These treatment-dependent differences in response latencies were not at the expense of other task performance measures however (Suppl. Fig 2H,K; Suppl. Fig 3D,H,P). Thus, although unilateral dDG dysfunction had some effect on task performance, there was no effect on disambiguation learning of similar spatial stimuli. Finally, to probe the spatial accuracy of choices, we quantified the response distributions for target (cued) and non-target (non-cued, i.e. blank) locations. The spatial distributions of nose pokes revealed no differences between the treatment conditions regardless of the separation distance and task phase (see Suppl. Table 1) for bilateral DREADD+ animals (*p*>0.05, Kolmogorov Smirnov test; Suppl. Fig 4A-B), unilateral DREADD+ animals (*p*>0.05, Kolmogorov Smirnov test; Suppl. Fig 4C-D) and control DREADD-animals (*p*>0.05, Kolmogorov Smirnov test; Suppl. Fig 4 E-F). These findings suggest that dDG inactivation did not impair the animal’s ability to locate the cued stimulus locations in space. Together, our findings indicate that CNO did not affect the animal’s ability to report its choices during the discrimination task, nor the ability to spatially locate the cued target locations, except for their ability to consecutively and correctly respond to the rewarded side.

To examine the possibility that the observed discrimination deficit in bilateral DREADD+ rats (Fig 2F) was caused by side effects of CNO being metabolized into clozapine on locomotion behaviour, we also quantified the movement speed of animals expressing DREADDs bilaterally during task engagement (Suppl. Fig 5A,C; see Suppl. Table 1). There was an interaction effect between separation and treatment on body speed during the acquisition phase (*F*(28)=5.49, *p*=0.027, GLMM), where animals moved significantly faster for CNO compared to VEH treatment during the acquisition phase (*W*=2.58, *p*=0.031, Wald test) and after the second reversal of Small sessions (*W*=3.27, *p*=0.0063, Wald test; Fig 2N; Suppl. Fig 5D). Interestingly, this task-phase and treatment-dependent effect on body speed was observed selectively when animals made correct (Suppl. Fig 5G) and leftward choices (Suppl. Fig 5K). In contrast, no effect of CNO on body speed was observed during Large sessions (Fig 2L; Suppl. Fig 5B), independent of the correctness of the trial (Suppl. Fig 5E-F) and of whether a response was made to the left or right side (Suppl. Fig 5I-J). Thus, the effect of CNO on body speed did not generalize across separation levels, suggesting that there was no general effect of CNO administration on locomotion activity.

### 3.2. Increased choice alternation underlies impaired discrimination learning during DG modulation

The dDG-dependent discrimination deficit observed during the acquisition phase (Fig 2F) was characterized by an increased number of trials needed to make consecutive correct choices to reach TTC, indicative of slower learning across trials. However, an alternative interpretation could be a deficit in the stability of choice behaviour due to animals being more uncertain when spatial discrimination is ambiguous.

To discern between these two possible interpretations, we set out to capture the learning dynamics across trials after DG modulation in more detail by quantifying the volatility of choice behaviour by assessing the tendency of animals to alternate or persevere in their choice. For this, we modelled the choice behaviour with two variants of a reinforcement learning (RL) model (Rescorla-Wagner, 1979; Sutton and Barto, 2018; Metha et al. 2020). The first RL model that we used (henceforth: ‘Simple RL model’) contained two parameters to characterize the learning dynamics: the learning rate (*alpha*; Fig 3D-E), which indicates how quickly animals adapt their choice across trials (i.e. reflecting the association strength between location and reward), and reward sensitivity (*beta*; Fig 3F-G), which indicates how sensitive animals are to differences in reward value. The second RL model (henceforth: ‘Perseverance RL model’; Metha et al. 2020) also included an additional parameter besides the learning rate and reward sensitivity: the perseverance rate (*delta*; Fig 3H-I), indicating the tendency of animals to persevere in their choice behaviour.

**Figure 3.**
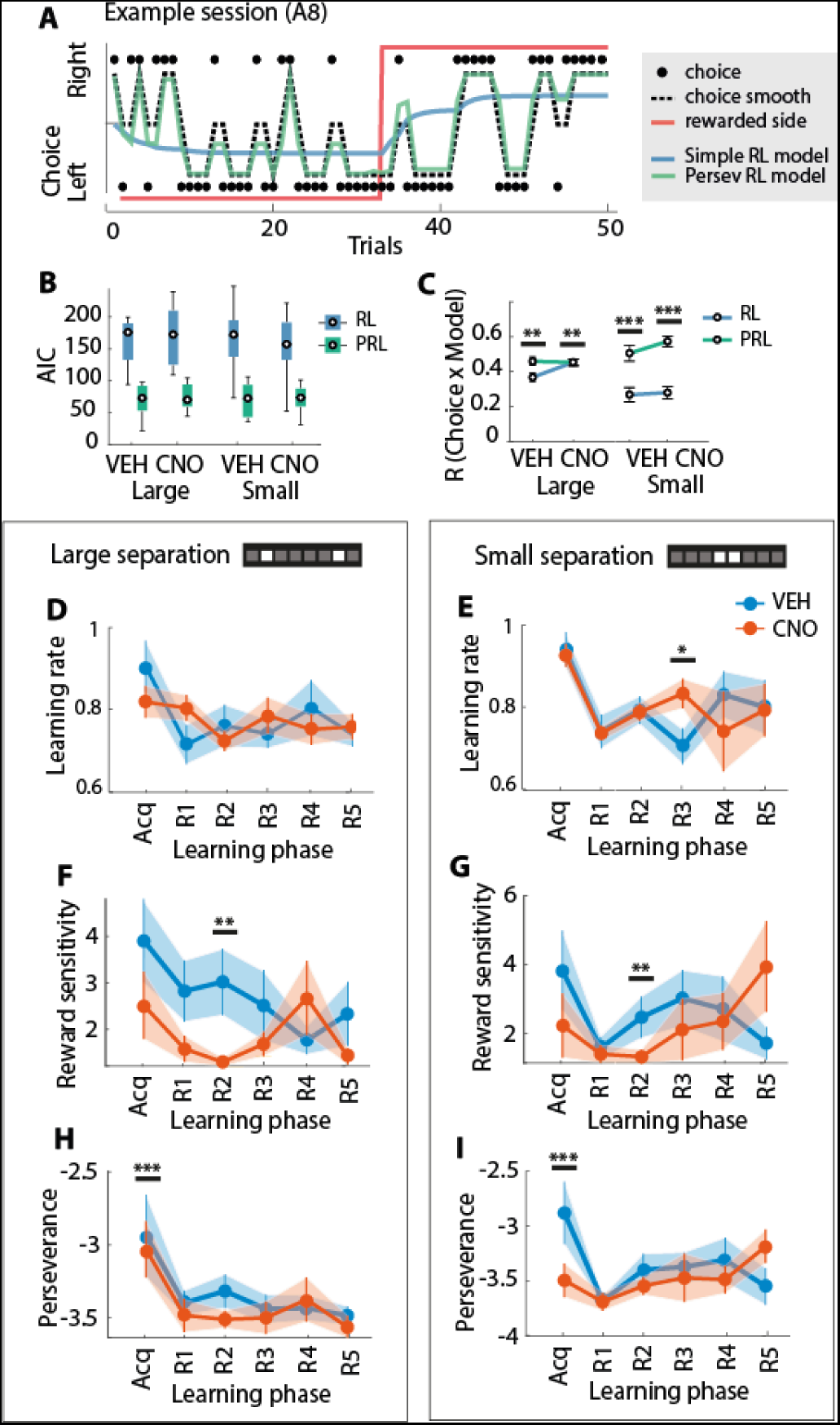
Increased choice switching associated with bilateral DG dependent discrimination deficit during acquisition learning of place reward associations. **A)** Representative example session of choice behaviour per trial (black dots) and smoothened across 4 trials (back dotted line) of bilateral DREADD+ animal 8 (A8) during a Small separation session with saline treatment. The choice trace is overlaid with the modelled behaviour from the Simple Reinforcement learning (RL) model (blue line) and the Perseverance RL (PRL) model (green line). The rewarded side is indicated with the red line ( reward side switches after the animal consecutive correct trials). All animals included in subsequent panels are from the bilateral DREADD+ cohort. **B)** Median Akaike Information Criterion (AIC) of the RL (blue square) and PRL model fits (green square) per treatment and separation condition. **C)** Per-session correlations (R) between the observed and modelled choices of each treatment and separation condition as fitted by the RL (blue circle) and PRL model (green circle). **D)** Mean learning rate for each learning phase with saline (VEH, blue line) or CNO (orange line) treatment for Large sessions. **E)** Same as D, but for Small sessions. **F,G)** Same as D) and E), respectively, but for the mean reward sensitivity. **H,I)** Same as D) and E), respectively, but for the mean perseverance rate. Boxplots indicate the interquartile range (IQR) of task variables pooled across animals (median ± IQR). Error bars indicate average task variables pooled across animals (mean± SEM). Significance is indicated by * p<0.05, ** p<0.01, *** p<0.001.

We fitted the RL models for each individual animal, task condition and learning phase on a trial-by-trial basis, which allowed us to obtain the parameter estimates separately for each rat per task condition and learning phase. The Perseverance RL model returned a lower Akaike Information Criterion (AIC) value compared to the simple RL model for every model fit for bilateral DREADD+ animals (Fig 3D), unilateral DREADD+ animals (Suppl. Fig 6A), and control DREADD-animals (Suppl. Fig 6C). Given that a lower AIC value indicates a significantly better model fit, these findings suggested that the variability in choice behaviour was better captured if we took the tendency of animals to persist in their choices into account. Moreover, the trial-by-trial estimates of the Perseverance RL model predictions correlated significantly stronger with the observed animals’ choice behaviour compared to the simple RL models (Fig 3C; Suppl. Fig 6B), except for the models fitted for control DREADD-animals (Suppl. Fig 6D). Our findings thus indicate that the Perseverance RL model is a better fit of choice behaviour compared to the Simple RL model.

We then compared the best fit parameter values across conditions for the Perseverance RL model for each DREADD expression group separately (see Suppl. Table 1). For bilateral DREADD+ animals, there was a main effect on learning rate not during the acquisition phase, but instead after the third reversal was made for Small separation sessions, where the learning rate was higher for CNO compared to vehicle treatment (*W*=2.67, *p*=0.022, Wald test; Fig 3E). In contrast, unilateral DREADD+ animals showed an interaction effect between treatment and separation on learning rate (*F*(9)=6.18, *p*=0.035, GLMM) after the fourth reversal was made for Large sessions only (Suppl. Fig 6E). The learning rate was, again, higher after CNO injections (*W*=3.17, *p*=0.023, Wald test). No effects of treatment were observed for the control DREADD-group (*p*>0.05, GLMM; Suppl. Fig 6G-H). This suggests that dDG dysfunction in at least one hemisphere modulated how rapidly animals adjusted their reward-driven choice behaviour during the reversal learning phase.

Interestingly, there was a main effect of treatment on the sensitivity to place-reward conditioning not during the acquisition phase, but after the second reversal for bilateral DREADD+ animals (*F*(36)=16.18, *p*=0.00028, GLMM), where reward sensitivity was significantly lower for CNO compared to VEH treatment for both Large (*W*=3.51, *p*=0.0012, Wald test, Fig 3F) and Small sessions (*W*=4.02, *p*=0.0006, Wald test; Fig 3G). Similarly, an interaction effect between separation and treatment was observed after the third reversal for unilateral DREADD+ animals (*F*(12)=366.35, *p*=2.33E-10), where reward sensitivity was lower after CNO injection compared to VEH for both Large (*W*=8.39, *p*=2.29E-04, Wald test; Suppl. Fig 6I) and Small sessions (*W*=38.35, *p*=0.00E-05, Wald test; Suppl. Fig 6J). No effect of treatment was observed on reward sensitivity for control DREADD-animals (*p*>0.05, GLMM, Suppl. Fig 6K-L). Our findings thus suggest that dDG dysfunction in at least one hemisphere reduced the sensitivity to place-reward conditioning during reversal learning, with no relation to separation distance.

Finally, there was a main effect of treatment on the perseverance rate during the acquisition phase for bilateral DREADD+ animals (*F*(40)=64.54, *p*=7.11E-10, GLMM), where the tendency to alternate across subsequent trials was higher after CNO injections compared to VEH for both Large (*W*=5.74, *p*=1.08E-04, Wald test; Fig 3H) and Small sessions (*W*=8.03, *p*=1.00E-09, Wald test; Fig 3I). Similarly, an interaction effect between separation and treatment was observed during acquisition learning for the unilateral DREADD+ cohort (*F*(20)=19.22, *p*=2.87E-04, GLMM), where the perseveration rate was lower for CNO compared to VEH treatment for Small sessions (*W*=2.48, *p*=0.044, Wald test; Suppl. Fig 6N). No effect of treatment was observed on the perseverance rate for the control DREADD-cohort (*p*>0.05, GLMM; Suppl. Fig 6O-P). Taken together, the Perseverance RL model revealed that the dDG-driven discrimination deficit observed during the acquisition phase of Small (Fig 2F) sessions was temporally associated with changes in alternation behaviours, but not in reward-driven choice behaviour or sensitivity to place reward conditioning.

## 4. Discussion

The main aim of the present study was to investigate the acute and causal role of the dorsal DG (dDG) in disambiguating similar spatial stimuli. Using inhibitory DREADDs, we showed that dysfunction of the dDG during a location discrimination task caused a discrimination deficit for similar locations during acquisition learning (Fig 2D,F). Our findings agree with the disambiguation function attributed to the DG, as probed behaviourally by previous studies lesion (Gilbert et al. 2001; Hunsaker and Kesner, 2008; Lee and Solivan, 2009; Morris et al. 2012;) and gene-knockout studies in rodents (McHugh et a. 2007; Kannangara et al. 2015; Yun et al. 2023). Importantly, we extend these prior studies by three novel findings. Firstly, we showed that the dDG was transiently engaged during discrimination learning by using reversible chemogenetic DREADDs instead of permanently perturbing dDG function with lesioning. Specifically, the task performance rapidly recovered to control levels after acquiring the initial ambiguous place-reward associations (Fig 2F). Secondly, we show that bilateral (Fig 2D,F), but not unilateral (Suppl. Fig 2K,N), modulation of the dDG induces a discrimination deficit for similar locations, suggesting that both hemispheres of the dDG are engaged during discrimination learning of similar locations. Lastly, our modelling study revealed that this initial deficit is not associated with changes in reward-driven behaviour (Fig 3E) nor sensitivity to place-reward conditioning (Fig 3G), but is instead associated with an increase in alternation behaviour (Fig 3I).

### 4.1. Transient bilateral modulation of the dorsal DG causes disambiguation deficits – chemogenetics versus permanent lesions

Our main finding that animals had difficulty disambiguating similar spatial stimuli when the DG function is perturbed agrees with previous lesion (Gilbert et al. 2001; Hunsaker and Kesner, 2008; Morris et al. 2012; Lee and Solivan, 2009) and gene knock-out studies (McHugh et a. 2007; Kannangara et al. 2015; Yun et al. 2023) in rodents. Although a direct comparison between studies is challenging due to task design differences, disambiguation deficits attributed to the DG have been more profound in previous studies compared to our study, showing that animals made more incorrect choices on average (Gilbert et al. 2001; Lee and Solivan, 2009; Oomen et al. 2015), while such an effect was absent in our study (Fig 2H,J). Instead, we observed a more subtle behavioural deficit, with animals responding more frequently incorrectly within a sequence of 9 out of 10 trials (see also Clelland et al. 2009; Morris et al. 2012) compared to control treatment sessions (Fig 2F). Moreover, we observed a DG-dependent discrimination deficit for 6 cm but not 30 cm separation in our study (Fig 2D,F), while previous deficits have been reported for a higher range of separation distances (15 to 60 cm, Gilbert et al. 2001; Lee and Solivan, 2009; 6 to 24 cm, Oomen et al. 2015). Our findings thus show, for the first time to our knowledge, that spatial disambiguation learning may be more sensitive to DG dysfunction after lesioning compared to transient chemogenetic modulation.

Although the spread of damage to the DG by neurotoxins such as colchicine and ibotenic acid has been typically reported to be confined within this subregion (Gilbert et al. 2001; Lee and Kesner 2003; Lee and Kesner, 2004; but see Jerman et al. 2005), the DG may be more sensitive to dysfunction after lesioning compared to chemogenetics due to off-target effects on downstream areas (Otchy et al. 2015; Vaidya et al. 2019). In such a case, the neural circuit supporting discrimination learning may not only concern the DG, but also its downstream areas such as the CA3 and CA1. Although similar off-target effects may occur acutely upon chemogenetic modulation in our study, we argue that lesioning may have caused differential effects due to compensation of affected functions via other areas within the functional circuit (Zelikowsky et al. 2013; Hong et al. 2018; reviewed in Vaidya et al. 2019). For instance, mice with extensive lesions in the dorsal hippocampus recovered their impaired contextual fear learning ability after two weeks, but only when the prefrontal cortex was left intact post-lesioning, suggesting that the latter area was functionally involved in the compensation process (Zelikowsky et al. 2013). Aside from the off-target and compensation effects, previous studies have also shown that lesioning the DG induces hyperactivity (Barone et al. 1992; Emerich et al. 1990). To examine whether there was a confounding effect of motor movement after chemogenetic targeting of the DG, we quantified the body speed during task engagement. Animals with bilateral dDG dysfunction tended to move faster during the acquisition phase and after the second reversal of Small sessions under CNO (Fig 2N; Suppl. Fig 5D), and specifically with correct (Suppl. Fig 5G) and left-ward trials (Suppl. Fig 5K). These findings suggest that there was no general effect of chemogenetics on movement during the task. We thus presumed that the hyperactivity observed during acquisition learning (Fig 2N) was caused by a temporary dysfunction of the DG due to their temporal association, which may cohere with the greater task difficulty for Small separation sessions under CNO (Fig 2F).

Although we intended to target the dDG bilaterally, some animals showed unilateral DREADD+ expression (Suppl. Fig 1B). In this latter group, we observed no chemogenetic effect on disambiguation learning when the dDG was targeted unilaterally (Suppl. Fig 2K,N; Suppl. Fig 3), suggesting that there is no lateralization of the spatial disambiguation function of the dDG. Indeed, although animals responded quicker after the fourth and fifth reversal of Small sessions under CNO (Suppl. Fig 3L), task performance remained unaffected during this reversal period (Suppl. Fig 2L; Suppl. Fig 3H). Previous lesion studies have either excluded or reported no animals with unilateral damage to the DG (Gilbert et al. 2001; Hunsaker and Kesner, 2008; Morris et al. 2012; Lee and Solivan, 2009; Oomen et al. 2015). Our study thus provides novel evidence that the spatial disambiguation function of the DG is not lateralized to one hemisphere. This contrasts with studies that have proposed a lateralization of spatial discrimination abilities in the hippocampus of humans (Miller et al. 2018) and rodents (Song et al. 2020). One possible explanation of the absence of an effect in unilateral DREADD+ animals (Suppl. Fig 2K,N) could be that the functionally intact hemisphere of the DG acutely compensated for a transient unilateral loss-of-function. Alternatively, our selective targeting of excitatory DG cells using DREADDs with a CaMKII promotor may have targeted a functional population of the DG that was not causally involved in the lateralization of such spatial discrimination abilities. Given that it is still unknown whether and how the DG contributes to the lateralization of spatial disambiguation functions, future studies differentiating between left and right DG function are still needed. Taken together, our study provides novel evidence that the acute engagement of the dDG in at least one hemisphere is needed to successfully discriminate similar spatial stimuli.

### 4.2. Spatial discrimination learning: working memory, place-outcome learning or spatial learning

To understand which aspects of spatial learning were affected by dDG dysfunction, we separately assessed discrimination learning during the acquisition phase and subsequent reversal phase (Fig 1C), as well as the learning dynamics underlying place-outcome learning using a validated reinforcement learning model (O’Reilly and den Ouden, 2015; Metha et al. 2020). The transient causal engagement of the dDG only during the acquisition phase (Fig 2D,F) suggests that reversal learning was left intact and validates previous indications that this type of learning is hippocampus independent (Jones and Mishkin, 1972; McAlonan and Brown, 2003; Boulougouris et al. 2007). Likewise, such a transient DG engagement renders it unlikely that the discrimination deficit can be explained as a general working memory or perceptual impairment, as such a deficit would persist across reversals. We observed no response bias for large or small separated cued locations, suggesting that spatial response learning based on an egocentric reference frame was also intact in these animals (e.g. the reward location is at the left side of the animal and not at its right). This validates earlier findings that this type of response learning is hippocampus independent and relies instead on the dorsal striatum (Packard and McGaugh, 1996; Featherstone and McDonald, 2004).

Instead, the selective, causal engagement of the dDG only concerned the initial learning of place-reward associations when the spatial separation was small, pointing to a deficit in fine-grained discrimination of rewarded versus nonrewarded locations (McDonald and White, 1995; Gilbert et al. 2001; Hunsaker and Kesner, 2008; McHugh et a. 2007; Morris et al. 2012; Lee and Solivan, 2009; Kannangara et al. 2015; Yun et al. 2023). Specifically, the impaired ability to respond consecutively correctly to the target location despite receiving reward hints towards a learning or spatial deficit; either a deficit in learning which place was associated with reward or, alternatively, a deficit in spatially separating nearby stimulus locations. We observed no effect of modulating the dDG on the pace of place-reward learning (Fig 3D-E) nor on the sensitivity of choice behaviour to differences in reward value during the acquisition phase regardless of the separation distance (Fig 3F-G). These findings suggest that the learning rate of place-outcome association learning is not affected by dDG dysfunction. Although animals did not make more incorrect choices on average during the acquisition phase (Fig 2H,J), they tended to respond less consistently across consecutive trials (Fig 2D,F) and alternated more frequently (Fig 3I). This suggests that, instead, dDG dysfunction causes a deterioration in the stability of choice behaviour. Given that volatile choice switching is typically associated with dysfunction in frontal cortices (Deserno et al. 2020), we consider it more likely that the discrimination deficit is attributed to a deficit in learning or maintaining correct place-reward representations during acquisition of place-outcome associations, i.e. knowing which location yields a reward. However, an alternative interpretation could be that animals had trouble spatially separating the stimulus locations. Nevertheless, the spatial distribution of blank pokes surrounding the target locations was not different between treatment sessions (Suppl. Fig 4A-B), suggesting that animals with dDG dysfunction did not respond less precisely when CNO was administered compared to control treatment sessions. Taken together, we propose that bilateral dDG dysfunction causes a deficit in the ability to encode place-reward associations or the fine-grained coding of place herein, although the two alternatives cannot be discriminated with the available data.

### 4.3. Neural mechanisms supporting the disambiguation of place-outcome representations

Our finding that the dDG is selectively engaged during discrimination of similar but not dissimilar spatial stimuli observed in our study supports the disambiguation function of the DG (Gilbert et al. 2001; Hunsaker and Kesner, 2008; McHugh et al. 2007; Morris et al. 2012; Lee and Solivan, 2009; Yun et al. 2023). Previous studies have suggested that the DG supports disambiguation learning via a pattern separation process, where neural representations of similar spatial stimuli are orthogonalized during memory formation (Marr et al. 1971; Treves and Rolls, 1994; Rolls and Kesner 2006; Clelland et al. 2009; reviewed in Yassa and Stark, 2011). Here, the orthogonalized encoding of fine-grained places, trajectories, and contexts by the DG is thought to be a crucial process to successfully discriminate between ambiguous contexts (Tolman, 1948; Leutgeb et al. 2007; reviewed in Lisman et al. 2017; van Dijk and Fenton, 2018). The question remains, however, how the DG would precisely contribute to the disambiguation of similar place-outcome representations.

Previous studies have shown that fine-grained places associated with reward outcomes are encoded by place cells in the hippocampus (Hollup et al. 2001; Hölscher et al. 2003; Lansink et al. 2012; Gauthier and Tank, 2018; reviewed in Sosa and Giocomo, 2021). For instance, the density and firing rate of place fields of CA1 cells was enhanced near goal locations (Hollup et al. 2001; Hölscher et al. 2003; Gauthier and Tank, 2018). Moreover, place fields in the CA1 were reduced in size at reward sites in a Y-maze, especially when reward-predicting cue lights were lit (Lansink et al. 2012), suggesting that the encoding of places becomes more fine-grained when they are associated with positive outcome. A recent study for example suggested that the DG is acutely engaged during the discrimination of similar but distinct contexts when the animal experiences a negative outcome (van Dijk and Fenton, 2018). This study agrees with our own observations that 1) the DG is acutely engaged during spatial discrimination in a 2) physically identical environment with re-located goal locations when 3) animals are actively acquiring place-outcome associations.

Interestingly, van Dijk and Fenton (2018) also observed that place cells co-fire more strongly with inhibitory interneurons in the DG during active discrimination. Interneurons in the hippocampus have been shown to enhance the spatial resolution of neighbouring place cells through synaptic inhibition (Rolotti et al. 2022). Thus, the co-firing of place cells with interneurons during active spatial discrimination may be a complementary mechanism with which the DG is able to better discriminate between fine-grained places, contexts, or trajectories. In turn, the dysfunction of the local circuit involving these subpopulations may explain why we observed a discrimination deficit specifically for the similar stimulus condition. Our histology revealed dense DREADD+ expressing cells in the hilar region (Fig 1F-G; Suppl. Figure 1A) containing mostly mossy cells (Senzai and Buszaki, 2017; Goodsmith et al. 2017). Mossy cells typically exhibit multiple place fields (Neunuebel and Knierim, 2012; Goodsmith et al. 2017) and are sensitive to subtle contextual changes (Nakazawa 2017; Senzai and Buszaki, 2017), suggesting that this subpopulation of the DG supports the disambiguation of spatial contexts. Moreover, hilar mossy cells have been shown to strongly drive granule cells (Jinde et al. 2012; Scharfman, 1990) that project onto the CA3 (reviewed in Amaral and Witter, 1989), which in turn is thought to be important for associative learning of places with objects and sensory stimuli (Morris et al. 2012). The behavioral effects that we observed in the present study could therefore have arisen due to a disruption of a DG-driven facilitation of the binding of motivationally salient events to exact places by downstream areas such as CA3, CA1-subiculum and ventral striatum.

### 4.4. The hippocampal-striatal-prefrontal system in spatial navigation and goal-directed decision making

In a broader context, the fine-grained discrimination between places, trajectories and environments is crucial for successful spatial navigation and goal-directed decision making. In support of this, the present study showed that animals with dDG dysfunction were less able to choose the correct goal location when the choice of spatial reward location was difficult. Previous studies have proposed that interactions between hippocampal, striatal, and prefrontal systems support spatial navigation and goal-directed decision making. Within this circuit, the hippocampus is thought to provide spatial coding signals and memorized information to the medial prefrontal cortex, while the ventral striatum – integrating signals from hippocampus and medial PFC – provides signals on reward expectancy and delivery to downstream areas modulating behaviour via direct connections to the prefrontal cortex (Alexander et al., 1986; Thierry et al. 2000; Voorn et al. 2004; Haber et al., 2006; Roberts et al., 2007). These hippocampal-prefrontal-striatal connections have been hypothesized to enable the prefrontal cortex to update behavioural task states and support flexible decision making (Brown et al., 2012, 2016; Brown and Stern, 2014; Ferbinteanu, 2016; Rusu and Pennartz 2020). This, in turn, suggests how the dDG dysfunction observed in our study may be associated with impaired decision making. Specifically, degraded spatial output from the hippocampus, relayed to the ventral striatum and prefrontal cortex, may have rendered the hippocampal-prefrontal-ventral striatal system unable to choose the correct goal location, despite being associated with reward. It remains to be investigated how the DG in particular contributes to this neural circuit supporting decision making and spatial navigation.

### 4.5 Conclusions

In conclusion, our study provides evidence for a mechanistic involvement of the dDG in acquisition learning of ambiguous place-reward associations. Future studies will have to uncover the circuit-level underpinnings of this role, in particular for what pertains the interplay between the dDG and other brain regions, both within and outside the hippocampal system.

## Acknowledgements

This study was supported by a ZonMW Off Road grant to C.A.O. and an Amsterdam Brain and Mind Project to C.S.L. We are grateful to Michel v.d. Oever (Vrije Universiteit Amsterdam) and Guus Smit (Vrije Universiteit Amsterdam) for their expertise input on work involving the DREADDs. We thank HelloBio.com for providing CNO, Alexander Mathis and Mackenzie Mathis (École Polytechnique Fédérale de Lausanne) for the DeepLabCut software tools provided for video tracking analysis, Hanneke den Ouden (Radboud University) and Jill O’Reilly (Oxford University) for providing the software for the reinforcement learning model.

## Contributions by authors

JL, ES and PA collected the behavioural data. JL analysed the behavioural data with feedback from UO and CP. JL and GH performed the surgeries. JL, ES and PA performed histology. JL wrote the manuscript with extensive feedback from UO and CP. CO designed the behavioural task. CL, CP and CO created the general study design.

## 6. Supplementary Figures

**Supplementary Figure 1 .**
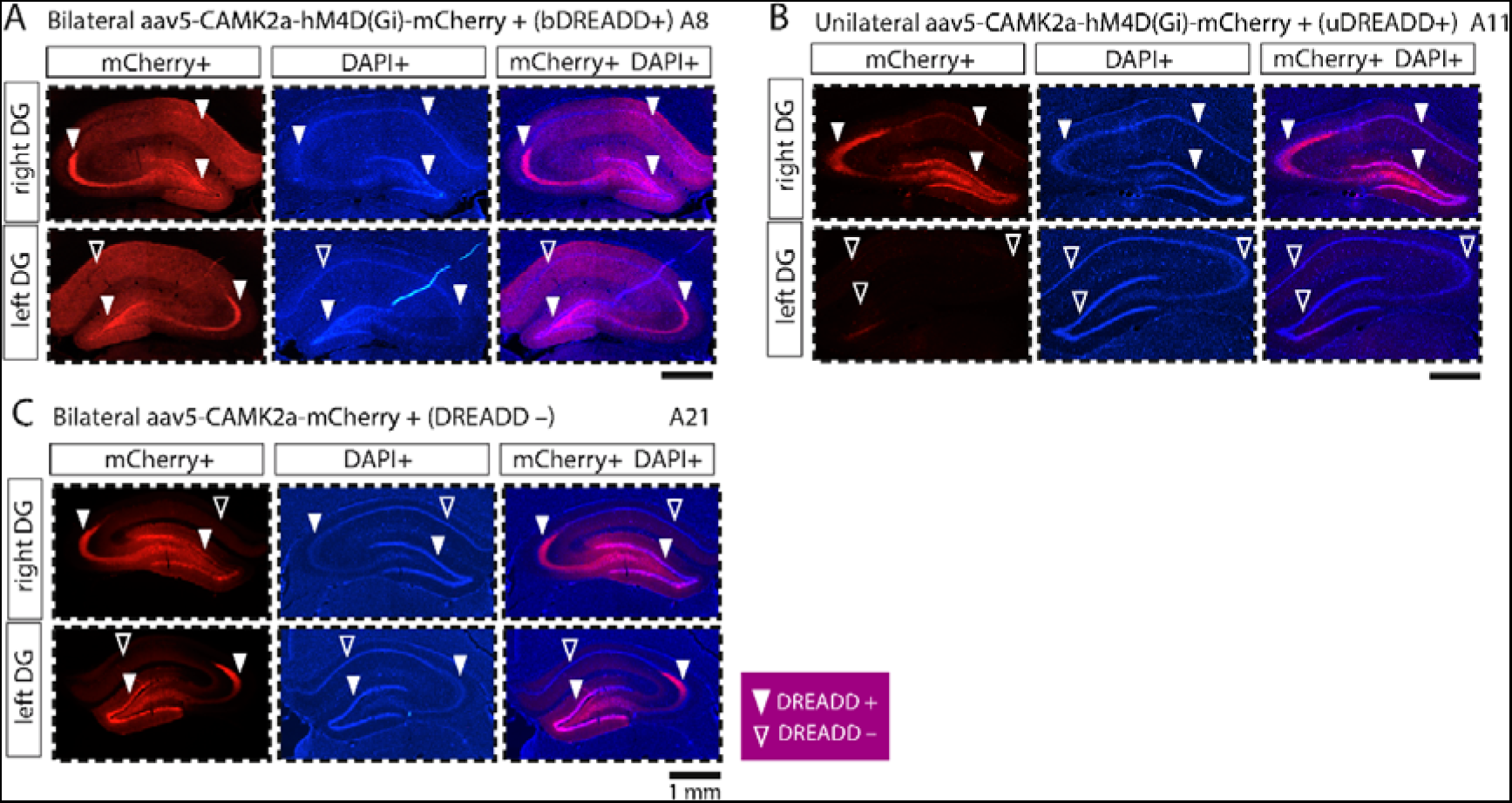
Histological verification of DREADD mCherry+ in the dorsal DG (dDG). **A)** Example animal 8 (A8) with bilateral DREADD mCherry expression in the dDG (left panel), DAPI expression (middle panel) and overlaid (right panel). **B)** Same as A), but for animal 11 (A11) with unilateral DREADD mCherry expression in the dDG.**C)** Same as B), but for animal 21 (A21) with bilateral mCherry expression with no DREADD vector (control virus) in the dDG. Solid triangles indicate subregions of the hippocampus expressing DREADDs. Outlined triangles indicate subregions of the hippocampus expressing no DREADDs.

**Supplementary Figure 2.**
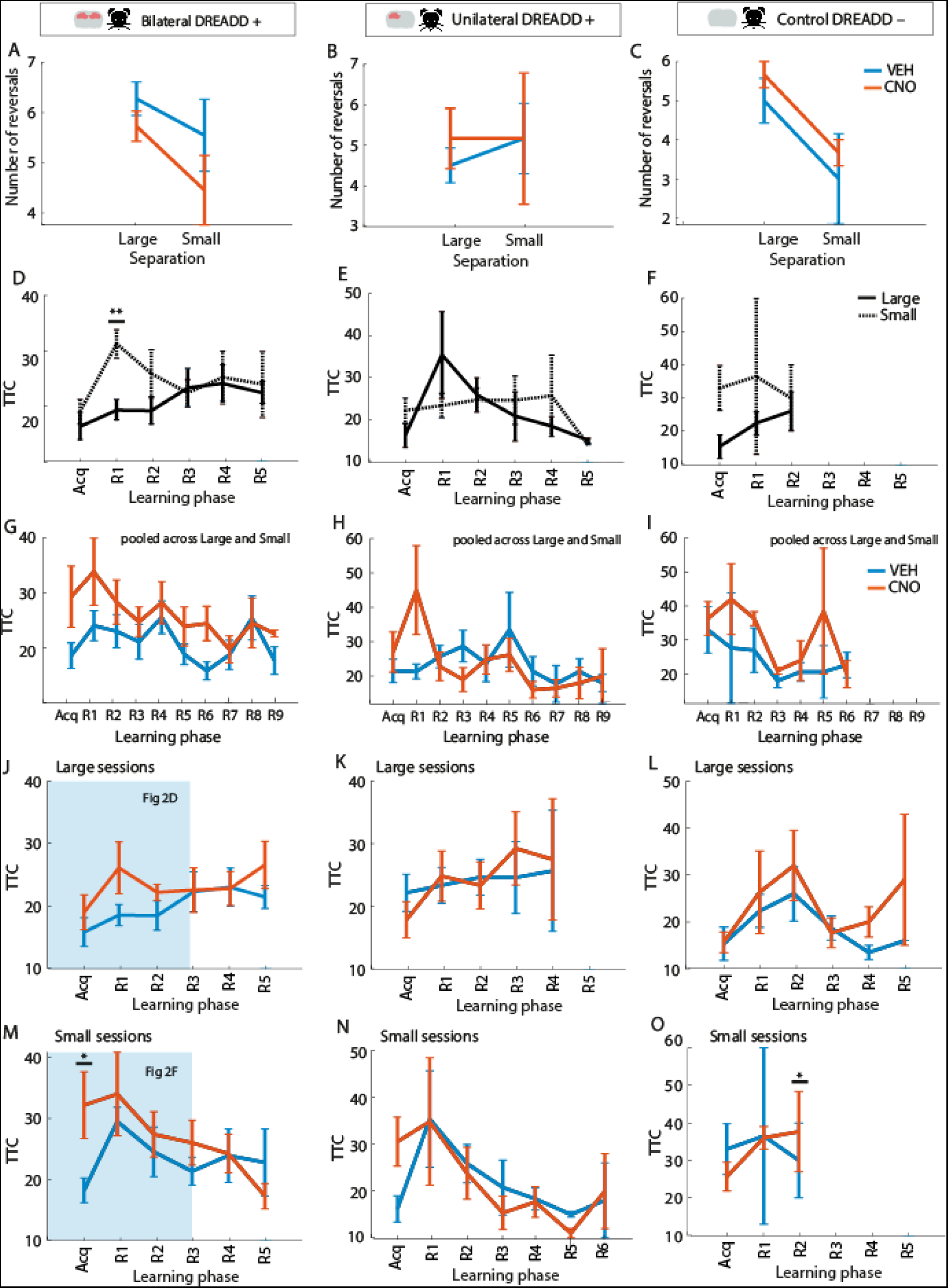
Quantification of number of reversals and trials needed to reach the reversal criterion for each treatment and separation condition across learning phases. **A)** Average number of reversals pooled across bilateral DREADD+ animals (N=11) for Large and Small sessions after treatment with saline (blue line) and CNO (orange line). **B)** Same as A), but for unilateral DREADD+ animals (N=6) **C)** Same as A), but for control DREADD-animals (N=3). **D)** Effect of separation: average trials needed to reach the reversal criterion (TTC) per learning phase for baseline (VEH) sessions for Large (solid black line) and Small (dotted black line) sessions for bilateral DREADD+ animals. **E)** Same as D), but for unilateral DREADD+ animals. **F)** Same as E), but for control DREADD-animals . **G)** Effect of treatment: average TTC per learning phase pooled across Large and Small sessions for VEH (blue line) and CNO (orange line) treatment for bilateral DREADD+ animals. **H)** Same as G), but for unilateral DREADD+ animals. **I)** Same as H), but for control DREADD-animals. **J)** TTC per learning phase for Large sessions after treatment with VEH (blue line) or CNO (orange line). **K)** Same as J), but for unilateral DREADD+ animals. **L)** Same as K), but for control DREADD-animals. **M)** TTC per learning phase for Small sessions after treatment with VEH (blue line) or CNO (orange line). **N)** Same as M), but for unilateral DREADD+ animals. **O)** Same as N), but for control DREADD-animals. Error bars indicate average task variables pooled across animals (mean± SEM). Significance is indicated by * p<0.05, ** p<0.01, *** p<0.001. For panels J) and M) the shaded regions indicate the plots already presented in Fig 2.

**Supplementary Figure 3.**
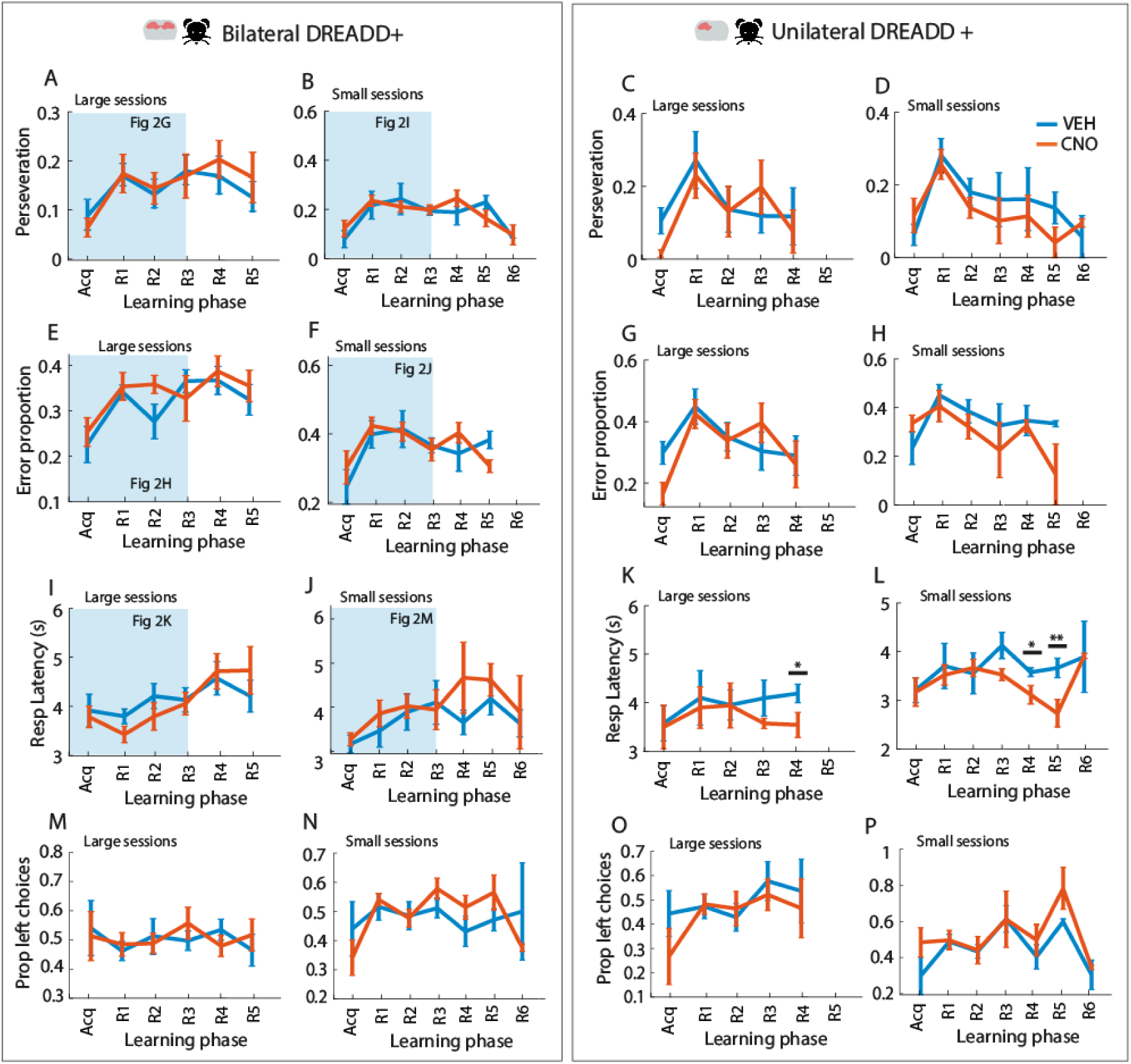
Behavioral task measures during location discrimination. **A)** Proportion of trials where bilateral DREADD+ animals (N=11) persisted in their choice relative to the previous trial for each learning phase for Large sessions only after saline (VEH, blue line) and CNO treatment (orange line). **B)** Same as A), but for Small sessions. **C-D)** Same as for A) to B), but for unilateral DREADD+ animals (N=6). **E)** Proportion of incorrect trials of bilateral DREADD+ animals for each learning phase for Large sessions. **F)** Same as E), but for Small sessions. **G-H)** Same as E) to F), but for unilateral DREADD+ animals. **I)** Response latency (in seconds, s) of bilateral DREADD+ animals for each learning phase after VEH or CNO treatment for Large sessions. **J)** Same as I), but for Small sessions**K**. **)-L)** Same as I) to J), but for unilateral DREADD+ animals. **M)** Proportion of leftward choice trials for bilateral DREADD+ animals for each learning phase after VEH or CNO treatment for Large sessions. **N)** Same as M), but for Small sessi**O**o)n**-P**s ). Same as M) to N), but for unilateral DREADD+ animals. Error bars indicate average task measures pooled across animals (mean± SEM). Significance is indicated by * p<0.05, ** p<0.01, *** p<0.001. Shaded regions in the plots related to bilateral DREADD+ animals indicate plots already shown in Fig 2.

**Supplementary Figure 4.**
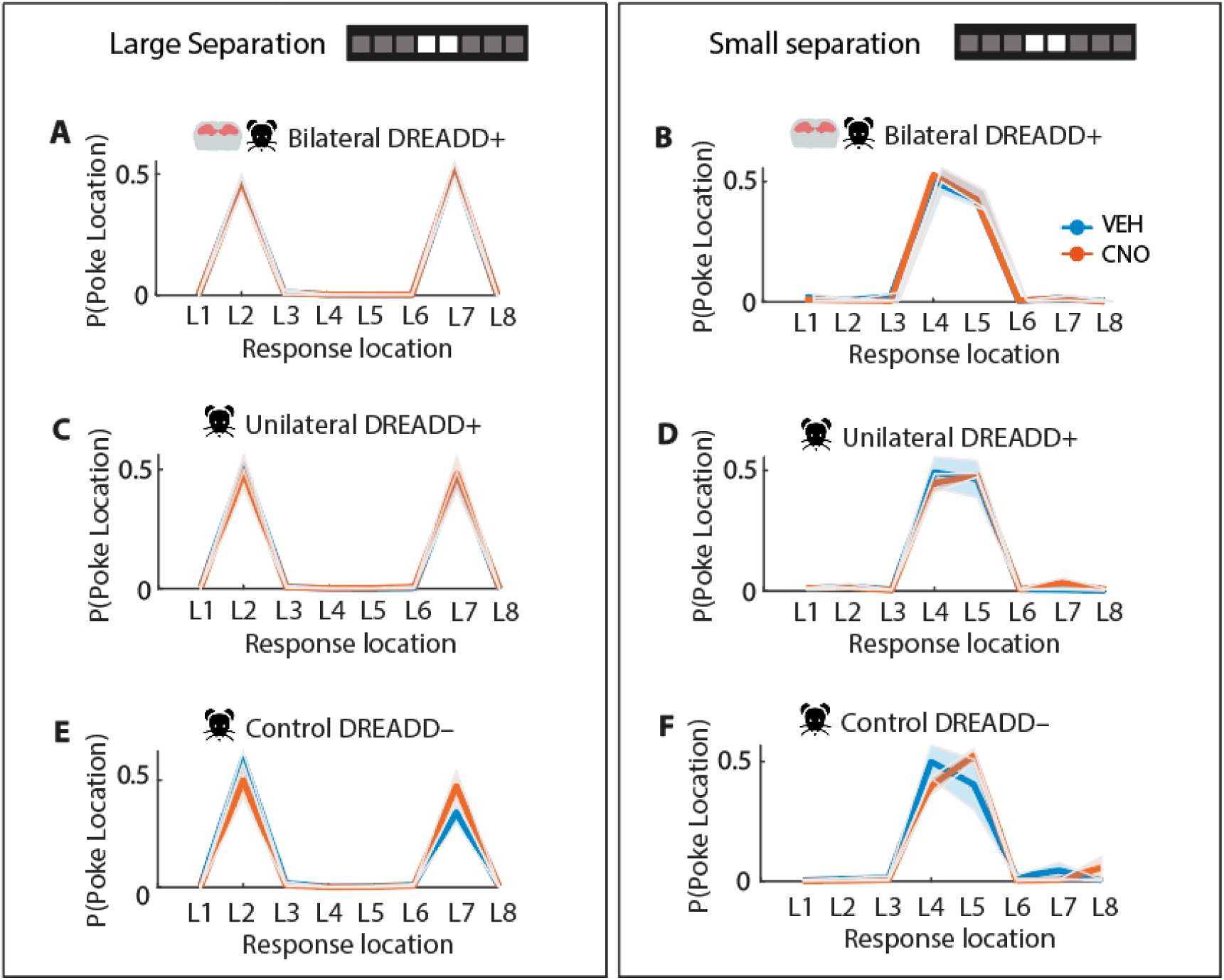
Spatial distribution of nose poke responses to location 1 to location 8. **A)** Poke probability of bilateral DREADD+ animals (N=11) to response locations 1 (L1) to 8 (L8) for Large sessions after saline (VEH, blue line) and CNO (orange line) treatment. The cued target and nontarget locations were alternated between L2 and L7 for subsequent reversals. **B)** Same as A), but for Small trials. The cued target and nontarget locations were alternated between L4 and L5 for subsequent reversals. **C-D)** Same as A) and B), but for unilateral DREADD+ animals (N=6). **E-F)** Same as C) and D), but for control DREADD-animals (N=3). Lines indicate poke probability (number of pokes at each location divided by the number of total pokes, average across animals. Shaded error bars indicate average poke probability pooled across animals (mean± SEM).

**Supplementary Figure 5.**
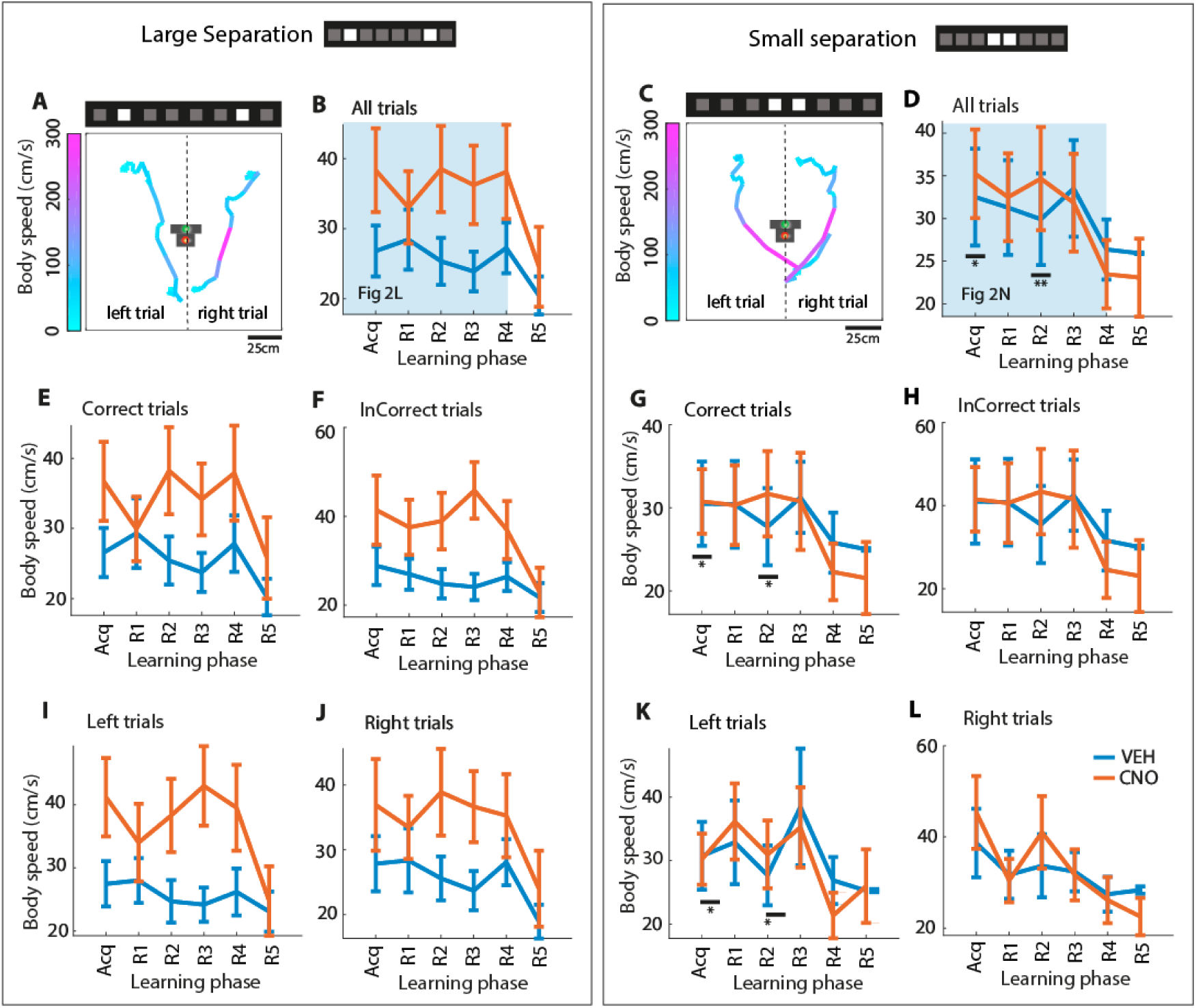
Body speed of animals during task engagement with DG modulation. **A)** Left: trajectory and body speed of a representative left trial for a Large VEH session of a bilateral DREADD+ animal (A9). Right: trajectory of a right trial for a Large CNO session of the same animal. Body speed (in cm/s) is indicated with colours scaled by the colour bar. **B)** Body speed of bilateral DREADD+ animals (N=8) for each learning phase for Large sessions after VEH and CNO treatment. All trials included. **C)** Same as A), but for a Small example session. **D)** Same as B), but for Small sessions. **E,G)** Body speed of bilateral DREADD+ animals for correct trials per treatment condition for Large and Small sessions, respectively. **F,H)** Body speed of bilateral DREADD+ animals for incorrect trials per treatment condition for Large and Small sessions, respectively. **I,K)** Body speed of bilateral DREADD+ animals for leftward trials per treatment condition for Large and Small sessions, respectively. **J,L)** Body speed of bilateral DREADD+ animalss for right ward trials per condition for Large and Small sessions, respectively. Error bars indicate average task variables pooled across animals (mean± SEM). Significance is indicated by * p<0.05, ** p<0.01, *** p<0.001. Shaded regions in panels B) and D) indicate plots already shown in Fig 2.

**Supplementary Figure 6.**
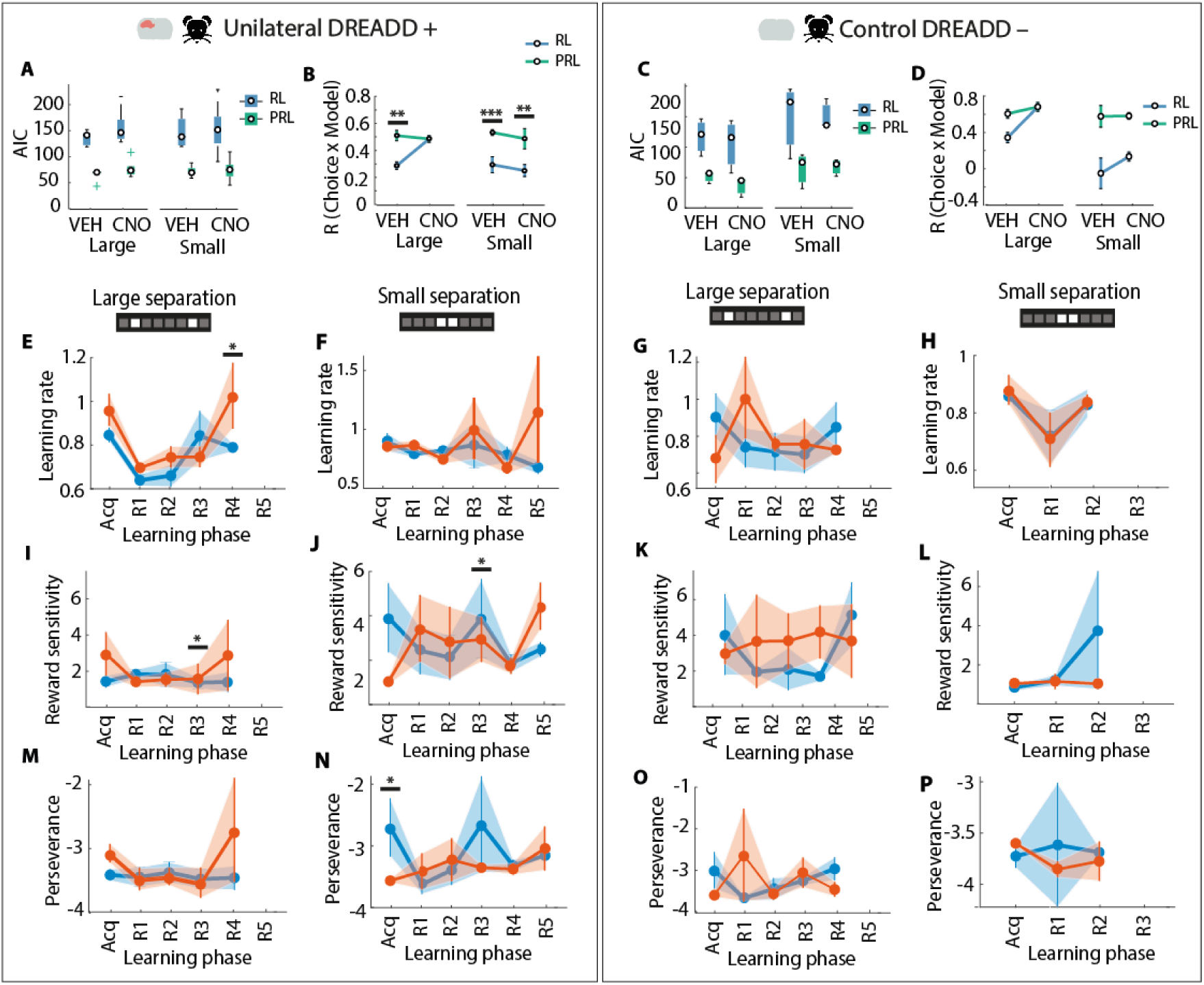
Choice behaviour fitted with Simple versus Perseverance reinforcement learning model for unilateral DREADD+ animals and control DREADD-animals. **A)** Median Akaike Information Criterion (AIC) of model fits and **B)** per-session correlations (R) between the observed and modelled choice of each treatment and separation condition as fitted by the Simple RL model (RL, blue square or circle) and Perseverance RL model (PRL, green square or circle) of unilateral DREADD+ animals (N=6). **C-D)** Same as A) and B), respectively, but or control DREADD-animals (N=3). **E)** Mean learning rate of unilateral DREADD+ animals for each learning phase with saline (VEH, blue square) or CNO (orange square) treatment for Large sessions and **F)** for Small sessions**G**. **-H)** Same as E) and F), but for control DREADD-animals. **I)** Mean reward sensitivity of unilateral DREADD+ animals per learning phase and treatment condition for Large and **J)** Small sessions. **K-L)** Same as I) and J), but for control DREADD-animals. **M)** Mean perseverance rate of unilateral DREADD+ animals per learning phase and treatment condition for Large sessions and **N)** Small sessions.**O-P)** Same as M) and N), but for control DREADD-animals. Boxplots indicate the interquartile range (IQR) of task variables pooled across animals (median ± IQR). Error bars indicate average task variables pooled across animals (mean± SEM). Significance is indicated by * p<0.05, ** p<0.01, *** p<0.001.

## References

Akaike, H. (1998) Information Theory and an Extension of the Maximum Likelihood Principle. In: Parzen, E., Tanabe, K. and Kitagawa, G., Eds., Selected Papers of Hirotugu Akaike, Springer Series in Statistics (Perspectives in Statistics), Springer, New York, 199–213.

Alexander GE, DeLong MR, Strick PL. Parallel organization of functionally segregated circuits linking basal ganglia and cortex. Annu Rev Neurosci. 1986;9:357–81. Doi: 10.1146/annurev.ne.09.030186.002041. PMID: 3085570.

Ally BA, Hussey EP, Ko PC, Molitor RJ. Pattern separation and pattern completion in Alzheimer’s disease: evidence of rapid forgetting in amnestic mild cognitive impairment. Hippocampus. 2013 Dec;23(12):1246–58. doi: 10.1002/hipo.22162. Epub 2013 Aug 14. PMID: 23804525; PMCID: PMC4310474.

Amaral DG, Witter MP. The three-dimensional organization of the hippocampal formation: a review of anatomical data. Neuroscience. 1989;31(3):571–91. doi: 10.1016/0306-4522(89)90424-7. PMID: 2687721.

Barone S Jr, Bonner M, Tandon P, McGinty JF, Tilson HA. The neurobiological effects of colchicine: modulation by nerve growth factor. Brain Res Bull. 1992 Feb;28(2):265–74. Doi: 10.1016/0361-9230(92)90188-4. PMID: 1596747.

Boulougouris V, Dalley JW, Robbins TW. Effects of orbitofrontal, infralimbic and prelimbic cortical lesions on serial spatial reversal learning in the rat. Behav Brain Res. 2007 May 16;179(2):219–28. doi: 10.1016/j.bbr.2007.02.005. Epub 2007 Feb 9. PMID: 17337305.

Brainard, D. H. (1997). The Psychophysics Toolbox. Spatial Vision, 10, 433–436. Doi: 10.1163/156856897X00357.

Brown, T. I., Ross, R. S., Tobyne, S. M., and Stern, C. E. (2012). Cooperative interactions between hippocampal and striatal systems support flexible navigation. Neuroimage 60, 1316–1330. Doi: 10.1016/j.neuroimage.2012.01.046

Brown, T. I., and Stern, C. E. (2014). Contributions of medial temporal lobe and striatal memory systems to learning and retrieving overlapping spatial memories. Cereb. Cortex 24, 1906–1922. Doi: 10.1093/cercor/bht041

Brown, T. I., Carr, V. A., LaRocque, K. F., Favila, S. E., Gordon, A. M., Bowles, B., et al. (2016). Prospective representation of navigational goals in the human hippocampus. Science 352, 1323–1326. Doi: 10.1126/science.aaf0784

Cavada, C., Compañy, T., Tejedor, J., Cruz-Rizzolo, R. J., and Reinoso-Suárez, F. (2000). The anatomical connections of the macaque monkey orbitofrontal cortex. A review. Cereb. Cortex 10, 220–242. Doi: 10.1093/cercor/10.3.220

Clelland, C.D., Choi, M., Romberg, C., Clemenson, G.D., Fragniere, A., Tyers, P., Jessberger, S., Saksida, L.M., Barker, R.A., Gage F.H., & Bussey, T.J. (2009). A functional role for adult hippocampal neurogenesis in spatial pattern separation. Science, 325, 210–213. Doi: 10.1126/science.1173215.

Coba, M.P., Komiyama, N.H., Nithianantharajah, J., Kopanitsa, M.V., Indersmitten, T., Skene, N.G., Tuck, E.J., Fricker, D.G., Elsegood, K.A., Stanford, L.E., Afinowi, N.O., Saksida, L.M., Bussey, T.J., O’Dell, T.J., & Grant, S.G (2012). TniK is required for postsynaptic and nuclear signaling pathways and cognitive function. Journal of Neuroscience, 32(40), 13987–13999. Doi: 10.1523/jneurosci.2433-12.2012.

Creer, D.J., Romberg, C., Saksida, L.M., van Praag H., & Bussey, T.J (2010). Running enhances spatial pattern separation in mice. Proceedings of the National Academy of Sciences of the United States of America, 107, 2367–2372. Doi: 10.1073/pnas.0911725107.

Deserno L, Boehme R, Mathys C, Katthagen T, Kaminski J, Stephan KE, Heinz A, Schlagenhauf F. Volatility Estimates Increase Choice Switching and Relate to Prefrontal Activity in Schizophrenia. Biol Psychiatry Cogn Neurosci Neuroimaging. 2020 Feb;5(2):173–183. doi: 10.1016/j.bpsc.2019.10.007. Epub 2019 Nov 5. PMID: 31937449.

Domi E, Xu L, Toivainen S, Nordeman A, Gobbo F, Venniro M, Shaham Y, Messing RO, Visser E, van den Oever MC, Holm L, Barbier E, Augier E, Heilig M. A neural substrate of compulsive alcohol use. Sci Adv. 2021 Aug 18;7(34):eabg9045. Doi: 10.1126/sciadv.abg9045. PMID: 34407947; PMCID: PMC8373126.

Dupret D, O’Neill J, Pleydell-Bouverie B, Csicsvari J. The reorganization and reactivation of hippocampal maps predict spatial memory performance. Nat Neurosci. 2010 Aug;13(8):995–1002. doi: 10.1038/nn.2599. Epub 2010 Jul 18. PMID: 20639874; PMCID: PMC2923061.

Eichenbaum H, Stewart C, Morris RG. Hippocampal representation in place learning. J Neurosci. 1990 Nov;10(11):3531–42. doi: 10.1523/JNEUROSCI.10-11-03531.1990. PMID: 2230943; PMCID: PMC6570096.

Eichenbaum H, Schoenbaum G, Young B, Bunsey M. Functional organization of the hippocampal memory system. Proc Natl Acad Sci U S A. 1996 Nov 26;93(24):13500–7. doi: 10.1073/pnas.93.24.13500. PMID: 8942963; PMCID: PMC33637.

Emerich DF, Walsh TJ. Hyperactivity following intradentate injection of colchicine: a role for dopamine systems in the nucleus accumbens. Pharmacol Biochem Behav. 1990 Sep;37(1):149–54. doi: 10.1016/0091-3057(90)90055-m. PMID: 2124711.

Featherstone RE, McDonald RJ. Dorsal striatum and stimulus-response learning: lesions of the dorsolateral, but not dorsomedial, striatum impair acquisition of a simple discrimination task. Behav Brain Res. 2004 Apr 2;150(1-2):15–23. doi: 10.1016/S0166-4328(03)00218-3. PMID: 15033275.

Ferbinteanu J, McDonald RJ. Dorsal/ventral hippocampus, fornix, and conditioned place preference. Hippocampus. 2001;11(2):187–200. doi: 10.1002/hipo.1036. PMID: 11345125.

Ferbinteanu, J. (2016). Contributions of hippocampus and striatum to memory-guided behavior depend on past experience. J. Neurosci. 36, 6459–6470. doi: 10.1523/JNEUROSCI.0840-16.2016

Gauthier JL, Tank DW. A Dedicated Population for Reward Coding in the Hippocampus. Neuron. 2018 Jul 11;99(1):179–193.e7. doi: 10.1016/j.neuron.2018.06.008. Epub 2018 Jun 28. PMID: 30008297; PMCID: PMC7023678.

Gilbert PE, Kesner RP, Lee I. Dissociating hippocampal subregions: double dissociation between dentate gyrus and CA1. Hippocampus. 2001;11(6):626–36. doi: 10.1002/hipo.1077. PMID: 11811656.

Gomez, J.M., Bonaventura, J., Lesniak, W., Mathews, W.B., Sysa-Shah, P., Rodriguez, L.A., Ellis, R.J., Richie, C.T., Harvey, B.K., Dannals, R.F., Pomper, M.G., Bonci, A., & Michaelides, M. (2017). Chemogenetics revealed: DREADD occupancy and activation via converted clozapine. Science, 357(6340), 503–507. doi: 10.1126/science.aan2475.

GoodSmith D, Chen X, Wang C, Kim SH, Song H, Burgalossi A, Christian KM, Knierim JJ. Spatial Representations of Granule Cells and Mossy Cells of the Dentate Gyrus. Neuron. 2017 Feb 8;93(3):677–690.e5. doi: 10.1016/j.neuron.2016.12.026. Epub 2017 Jan 26. PMID: 28132828; PMCID: PMC5300955.

Haber, S. N., Kim, K. S., Mailly, P., and Calzavara, R. (2006). Reward-related cortical inputs define a large striatal region in primates that interface with associative cortical connections, providing a substrate for incentive-based learning. J. Neurosci. 26, 8368–8376. doi: 10.1523/JNEUROSCI.0271-06.2006

Hainmueller T, Bartos M. Parallel emergence of stable and dynamic memory engrams in the hippocampus. Nature. 2018 Jun;558(7709):292–296. doi: 10.1038/s41586-018-0191-2. Epub 2018 Jun 6. PMID: 29875406; PMCID: PMC7115829.

Hampton RR, Murray EA (2004) Selective hippocampal damage in rhesus monkeys impairs spatial memory in an open-field test. Hippocampus 14:808–818.

Hollup SA, Molden S, Donnett JG, Moser MB, Moser EI. Accumulation of hippocampal place fields at the goal location in an annular watermaze task. J Neurosci. 2001 Mar 1;21(5):1635–44. doi: 10.1523/JNEUROSCI.21-05-01635.2001. PMID: 11222654; PMCID: PMC6762966.

Hölscher C, Jacob W, Mallot HA. Reward modulates neuronal activity in the hippocampus of the rat. Behav Brain Res. 2003 Jun 16;142(1-2):181–91. doi: 10.1016/s0166-4328(02)00422-9. PMID: 12798280.

Hong YK, Lacefield CO, Rodgers CC, Bruno RM. Sensation, movement and learning in the absence of barrel cortex. Nature. 2018 Sep;561(7724):542–546. doi: 10.1038/s41586-018-0527-y. Epub 2018 Sep 17. PMID: 30224746; PMCID: PMC6173956.

Hsiang HL, Epp JR, van den Oever MC, Yan C, Rashid AJ, Insel N, Ye L, Niibori Y, Deisseroth K, Frankland PW, Josselyn SA. Manipulating a “cocaine engram” in mice. J Neurosci. 2014 Oct 15;34(42):14115–27. doi: 10.1523/JNEUROSCI.3327-14.2014. PMID: 25319707; PMCID: PMC6705287.

Hunsaker MR, Kesner RP. Evaluating the differential roles of the dorsal dentate gyrus, dorsal CA3, and dorsal CA1 during a temporal ordering for spatial locations task. Hippocampus. 2008;18(9):955–64. doi: 10.1002/hipo.20455. PMID: 18493930; PMCID: PMC2570230.

Hunsaker MR, Kesner RP. The operation of pattern separation and pattern completion processes associated with different attributes or domains of memory. Neurosci Biobehav Rev. 2013 Jan;37(1):36–58. doi: 10.1016/j.neubiorev.2012.09.014. Epub 2012 Oct 5. PMID: 23043857.

Ito R, Robbins TW, Pennartz CM, Everitt BJ. Functional interaction between the hippocampus and nucleus accumbens shell is necessary for the acquisition of appetitive spatial context conditioning. J Neurosci. 2008 Jul 2;28(27):6950–9. doi: 10.1523/JNEUROSCI.1615-08.2008. PMID: 18596169; PMCID: PMC3844800.

Jerman TS, Kesner RP, Lee I, Berman RF. Patterns of hippocampal cell loss based on subregional lesions of the hippocampus. Brain Res. 2005 Dec 14;1065(1-2):1–7. doi: 10.1016/j.brainres.2005.09.062. Epub 2005 Nov 22. PMID: 16307731.

Jinde S, Zsiros V, Jiang Z, Nakao K, Pickel J, Kohno K, Belforte JE, Nakazawa K. Hilar mossy cell degeneration causes transient dentate granule cell hyperexcitability and impaired pattern separation. Neuron. 2012 Dec 20;76(6):1189–200. doi: 10.1016/j.neuron.2012.10.036. PMID: 23259953; PMCID: PMC3530172.

Jones B, Mishkin M. Limbic lesions and the problem of stimulus--reinforcement associations. Exp Neurol. 1972 Aug;36(2):362–77. doi: 10.1016/0014-4886(72)90030-1. PMID: 4626489.

Kannangara TS, Eadie BD, Bostrom CA, Morch K, Brocardo PS, Christie BR. GluN2A-/- Mice Lack Bidirectional Synaptic Plasticity in the Dentate Gyrus and Perform Poorly on Spatial Pattern Separation Tasks. Cereb Cortex. 2015 Aug;25(8):2102–13. doi: 10.1093/cercor/bhu017. Epub 2014 Feb 18. PMID: 24554729; PMCID: PMC4494024.

Khamassi, M., and Humphries, M. D. (2012). Integrating cortico-limbic-basal ganglia architectures for learning model-based and model-free navigation strategies. Front. Behav. Neurosci. 6:79. doi: 10.3389/fnbeh.2012.00079

Kirk, R. A., Redmon, S. N., & Kesner, R. P. (2017). The ventral dentate gyrus mediates pattern separation for reward value. Behavioral Neuroscience, 131(1), 42–45. 10.1037/bne0000172

Knierim JJ, Neunuebel JP. Tracking the flow of hippocampal computation: Pattern separation, pattern completion, and attractor dynamics. Neurobiol Learn Mem. 2016 Mar;129:38–49. doi: 10.1016/j.nlm.2015.10.008. Epub 2015 Oct 26. PMID: 26514299; PMCID: PMC4792674.

Laczó M, Lerch O, Martinkovic L, Kalinova J, Markova H, Vyhnalek M, Hort J, Laczó J. Spatial Pattern Separation Testing Differentiates Alzheimer’s Disease Biomarker-Positive and Biomarker-Negative Older Adults With Amnestic Mild Cognitive Impairment. Front Aging Neurosci. 2021 Nov 26;13:774600. doi: 10.3389/fnagi.2021.774600. PMID: 34899277; PMCID: PMC8662816.

Lansink CS, Goltstein PM, Lankelma JV, McNaughton BL, Pennartz CM. Hippocampus leads ventral striatum in replay of place-reward information. PLoS Biol. 2009 Aug;7(8):e1000173. doi: 10.1371/journal.pbio.1000173. Epub 2009 Aug 18. PMID: 19688032; PMCID: PMC2717326.

Lansink CS, Jackson JC, Lankelma JV, Ito R, Robbins TW, Everitt BJ, Pennartz CM. Reward cues in space: commonalities and differences in neural coding by hippocampal and ventral striatal ensembles. J Neurosci. 2012 Sep 5;32(36):12444–59. doi: 10.1523/JNEUROSCI.0593-12.2012. PMID: 22956836; PMCID: PMC3492752.

Latuske P, Kornienko O, Kohler L, Allen K. Hippocampal Remapping and Its Entorhinal Origin. Front Behav Neurosci. 2018 Jan 4;11:253. doi: 10.3389/fnbeh.2017.00253. PMID: 29354038; PMCID: PMC5758554.

Lavenex PB, Amaral DG, Lavenex P. Hippocampal lesion prevents spatial relational learning in adult macaque monkeys. J Neurosci. 2006 Apr 26;26(17):4546–58. doi: 10.1523/JNEUROSCI.5412-05.2006. PMID: 16641234; PMCID: PMC6674053.

Leal SL, Yassa MA. Integrating new findings and examining clinical applications of pattern separation. Nat Neurosci. 2018 Feb;21(2):163–173. doi: 10.1038/s41593-017-0065-1. Epub 2018 Jan 25. PMID: 29371654; PMCID: PMC5898810.

Lee JW, Kim WR, Sun W, Jung MW. Role of dentate gyrus in aligning internal spatial map to external landmark. Learn Mem. 2009 Aug 25;16(9):530–6. doi: 10.1101/lm.1483709. PMID: 19706836.

Lee JW, Jung MW. Separation or binding? Role of the dentate gyrus in hippocampal mnemonic processing. Neurosci Biobehav Rev. 2017;75:183–194. [PubMed] [Google Scholar] [Ref list]

Lee I, Kesner RP. Differential roles of dorsal hippocampal subregions in spatial working memory with short versus intermediate delay. Behav Neurosci. 2003 Oct;117(5):1044–53. doi: 10.1037/0735-7044.117.5.1044. PMID: 14570553.

Lee I, Kesner RP. Encoding versus retrieval of spatial memory: double dissociation between the dentate gyrus and the perforant path inputs into CA3 in the dorsal hippocampus. Hippocampus. 2004;14(1):66–76. doi: 10.1002/hipo.10167. PMID: 15058484.

Lee I, & Solivan F. Dentate gyrus is necessary for disambiguating similar object-place representations. Learn Mem. 2010 Apr 26;17(5):252–8. doi: 10.1101/lm.1678210. PMID: 20421312; PMCID: PMC2862407.

Lee CH, Lee I. Impairment of Pattern Separation of Ambiguous Scenes by Single Units in the CA3 in the Absence of the Dentate Gyrus. J Neurosci. 2020 Apr 29;40(18):3576–3590. doi: 10.1523/JNEUROSCI.2596-19.2020. Epub 2020 Mar 31. PMID: 32234778; PMCID: PMC7189760.

Lee H, Stirnberg R, Wu S, Wang X, Stöcker T, Jung S, Montag C, Axmacher N. Genetic Alzheimer’s Disease Risk Affects the Neural Mechanisms of Pattern Separation in Hippocampal Subfields. Curr Biol. 2020 Nov 2;30(21):4201–4212.e3. doi: 10.1016/j.cub.2020.08.042. Epub 2020 Sep 10. PMID: 32916120.

Lesuis SL, Brosens N, Immerzeel N, van der Loo RJ, Mitrić M, Bielefeld P, Fitzsimons CP, Lucassen PJ, Kushner SA, van den Oever MC, Krugers HJ. Glucocorticoids Promote Fear Generalization by Increasing the Size of a Dentate Gyrus Engram Cell Population. Biol Psychiatry. 2021 Oct 1;90(7):494–504. doi: 10.1016/j.biopsych.2021.04.010. Epub 2021 Apr 24. PMID: 34503674.

Leutgeb JK, Leutgeb S, Moser MB, Moser EI. Pattern separation in the dentate gyrus and CA3 of the hippocampus. Science. 2007 Feb 16;315(5814):961–6. doi: 10.1126/science.1135801. PMID: 17303747.

Lisman J, Buzsáki G, Eichenbaum H, Nadel L, Ranganath C, Redish AD. Viewpoints: how the hippocampus contributes to memory, navigation and cognition. Nat Neurosci. 2017 Oct 26;20(11):1434–1447. doi: 10.1038/nn.4661. Erratum in: Nat Neurosci. 2017 Dec 20;: PMID: 29073641; PMCID: PMC5943637.

Manvich DF, Webster KA, Foster SL, Farrell MS, Ritchie JC, Porter JH, Weinshenker D. The DREADD agonist clozapine N-oxide (CNO) is reverse-metabolized to clozapine and produces clozapine-like interoceptive stimulus effects in rats and mice. Sci Rep. 2018 Mar 1;8(1):3840. doi: 10.1038/s41598-018-22116-z. PMID: 29497149; PMCID: PMC5832819.

Marr D. Simple memory: a theory for archicortex. Philos Trans R Soc Lond B Biol Sci. 1971 Jul 1;262(841):23–81. doi: 10.1098/rstb.1971.0078. PMID: 4399412.

Mathis, A., Mamidanna, P., Cury, K.M., Abe, T., Murthy, V.N., Mathis, M.W., & Bethge, M. (2018). DeepLabCut: markerless pose estimation of user-defined body parts with deep learning. Nature Neuroscience, 21, 1281–1289. doi: 10.1038/s41593-018-0209-y.

McAlonan K., Brown V.J. Orbital prefrontal cortex mediates reversal learning and not attentional set shifting in the rat. Behav Brain Res. 2003;146:97–103. [PubMed] [Google Scholar]

McDonald RJ, White NM. Hippocampal and nonhippocampal contributions to place learning in rats. Behav Neurosci. 1995 Aug;109(4):579–93. doi: 10.1037//0735-7044.109.4.579. PMID: 7576202.

McHugh TJ, Jones MW, Quinn JJ, Balthasar N, Coppari R, Elmquist JK, Lowell BB, Fanselow MS, Wilson MA, Tonegawa S. Dentate gyrus NMDA receptors mediate rapid pattern separation in the hippocampal network. Science. 2007 Jul 6;317(5834):94–9. doi: 10.1126/science.1140263. Epub 2007 Jun 7. PMID: 17556551.

MacLaren DA, Browne RW, Shaw JK, Krishnan Radhakrishnan S, Khare P, España RA, Clark SD. Clozapine N-Oxide Administration Produces Behavioral Effects in Long-Evans Rats: Implications for Designing DREADD Experiments. eNeuro. 2016 Nov 1;3(5):ENEURO.0219-16.2016. doi: 10.1523/ENEURO.0219-16.2016. PMID: 27822508; PMCID: PMC5089539.

McNaughton BL, Barnes CA, Gerrard JL, Gothard K, Jung MW, Knierim JJ, Kudrimoti H, Qin Y, Skaggs WE, Suster M, Weaver KL. Deciphering the hippocampal polyglot: the hippocampus as a path integration system. J Exp Biol. 1996 Jan;199(Pt 1):173–85. doi: 10.1242/jeb.199.1.173. PMID: 8576689.

McTighe, S.M., Mar, A.C., Romberg, C., Bussey, T.J., & Saksida, L.M. (2009). A new touchscreen test of pattern separation: effect of hippocampal lesions. Neuroreport, 20, 881–885. doi: 10.1097/WNR.0b013e32832c5eb2.

Metha, J.A., Brian, M.L., Oberrauch, S., Barnes, S.A., Featherby, T.J., Bossaerts, P., Murawski, C., Hoyer, D., & Jacobson, L.H. (2020). Separating probability and reversal learning in a novel probabilistic reversal learning task for mice. Frontiers in Behavioral Neuroscience, 13(270), 1–9. doi: 10.3389/fnbeh.2019.00270.

Middleton, F. A., and Strick, P. L. (2002). Basal-ganglia ‘projections’ to the prefrontal cortex of the primate. Cereb. Cortex 12, 926–935. doi: 10.1093/cercor/12.9.926

Miller J, Watrous AJ, Tsitsiklis M, Lee SA, Sheth SA, Schevon CA, Smith EH, Sperling MR, Sharan A, Asadi-Pooya AA, Worrell GA, Meisenhelter S, Inman CS, Davis KA, Lega B, Wanda PA, Das SR, Stein JM, Gorniak R, Jacobs J. Lateralized hippocampal oscillations underlie distinct aspects of human spatial memory and navigation. Nat Commun. 2018 Jun 21;9(1):2423. doi: 10.1038/s41467-018-04847-9. PMID: 29930307; PMCID: PMC6013427.

Morris AM, Churchwell JC, Kesner RP, Gilbert PE. Selective lesions of the dentate gyrus produce disruptions in place learning for adjacent spatial locations. Neurobiol Learn Mem. 2012 Mar;97(3):326–31. doi: 10.1016/j.nlm.2012.02.005. Epub 2012 Feb 27. PMID: 22390856; PMCID: PMC4089983.

Nakazawa K. Dentate Mossy Cell and Pattern Separation. Neuron. 2017 Feb 8;93(3):465–467. doi: 10.1016/j.neuron.2017.01.021. Erratum in: Neuron. 2017 Mar 8;93(5):1236. PMID: 28182899.

Neunuebel JP, Knierim JJ. Spatial firing correlates of physiologically distinct cell types of the rat dentate gyrus. J Neurosci. 2012 Mar 14;32(11):3848–58. doi: 10.1523/JNEUROSCI.6038-11.2012. PMID: 22423105; PMCID: PMC3321836.

Olton DS, Werz MA. Hippocampal function and behavior: spatial discrimination and response inhibition. Physiol Behav. 1978 May;20(5):597–605. doi: 10.1016/0031-9384(78)90252-4. PMID: 684094.

O’Keefe J, Dostrovsky J. The hippocampus as a spatial map. Preliminary evidence from unit activity in the freely-moving rat. Brain Res. 1971 Nov;34(1):171–5. doi: 10.1016/0006-8993(71)90358-1. PMID: 5124915.

O’Keefe J. Place units in the hippocampus of the freely moving rat. Exp Neurol. 1976 Apr;51(1):78–109. doi: 10.1016/0014-4886(76)90055-8. PMID: 1261644.

O’Keefe, J; Nadel, L; (1978) The Hippocampus as a Cognitive Map. Oxford University Press: Oxford, UK

Oomen CA, Hvoslef-Eide M, Heath CJ, Mar AC, Horner AE, Bussey TJ, Saksida LM. The touchscreen operant platform for testing working memory and pattern separation in rats and mice. Nat Protoc. 2013 Oct;8(10):2006–21. doi: 10.1038/nprot.2013.124. Epub 2013 Sep 19. PMID: 24051961; PMCID: PMC3982138.

Oomen CA, Hvoslef-Eide M, Kofink D, Preusser F, Mar AC, Saksida LM, Bussey TJ. A novel 2- and 3-choice touchscreen-based continuous trial-unique nonmatching-to-location task (cTUNL) sensitive to functional differences between dentate gyrus and CA3 subregions of the hippocampus. Psychopharmacology (Berl). 2015 Nov;232(21-22):3921–3933. doi: 10.1007/s00213-015-4019-6. Epub 2015 Jul 30. PMID: 26220610; PMCID: PMC4976805.

O’Reilly and den Ouden, 2015. http://www.hannekedenouden.ruhosting.nl/RLtutorial/Instructions.html

Otchy TM, Wolff SB, Rhee JY, Pehlevan C, Kawai R, Kempf A, Gobes SM, Ölveczky BP. Acute off-target effects of neural circuit manipulations. Nature. 2015 Dec 17;528(7582):358–63. doi: 10.1038/nature16442. Epub 2015 Dec 9. PMID: 26649821.

Packard, M. G., and McGaugh, J. L. (1996). Inactivation of hippocampus or caudate nucleus with lidocaine differentially affects expression of place and response learning. Neurobiol. Learn. Mem. 65, 65–72. doi: 10.1006/nlme.1996.0007

Parizkova M, Lerch O, Andel R, Kalinova J, Markova H, Vyhnalek M, Hort J, Laczó J. Spatial Pattern Separation in Early Alzheimer’s Disease. J Alzheimers Dis. 2020;76(1):121–138. doi: 10.3233/JAD-200093. PMID: 32444544.

Rescorla, R.A., & Wagner, A.R. (1972). A theory of Pavlovian conditioning: variations in the effectiveness of reinforcement and nonreinforcement. From: Black, A.H. & Prokasy W.F. Classical conditioning II, page 64–99. New York, NY: Appleton-Century-Crofts.

Roberts, A. C., Tomic, D. L., Parkinson, C. H., Roeling, T. A., Cutter, D. J., Robbins, T. W., et al. (2007). Forebrain connectivity of the prefrontal cortex in the marmoset monkey (Callithrix jacchus): an anterograde and retrograde tract-tracing study. J. Comp. Neurol. 502, 86–112. doi: 10.1002/cne.21300

Rolls, E. T. (1996). A theory of hippocampal function in memory. Hippocampus, 6(6), 601–620. 10.1002/(SICI)1098-1063(1996)6:6<601::AID-HIPO5>3.0.CO;2-J

Rolls ET, Kesner RP. A computational theory of hippocampal function, and empirical tests of the theory. Prog Neurobiol. 2006 May;79(1):1–48. doi: 10.1016/j.pneurobio.2006.04.005. Epub 2006 Jun 14. PMID: 16781044.

Rolotti SV, Ahmed MS, Szoboszlay M, Geiller T, Negrean A, Blockus H, Gonzalez KC, Sparks FT, Solis Canales AS, Tuttman AL, Peterka DS, Zemelman BV, Polleux F, Losonczy A. Local feedback inhibition tightly controls rapid formation of hippocampal place fields. Neuron. 2022 Mar 2;110(5):783–794.e6. doi: 10.1016/j.neuron.2021.12.003. Epub 2022 Jan 5. PMID: 34990571; PMCID: PMC8897257.

Roth, B.L. (2016). DREADDs for neuroscientists. Neuron, 89(4), 683–694. doi: 10.1016/j.neuron.2016.01.040

Rusu SI, Pennartz CMA. Learning, memory and consolidation mechanisms for behavioral control in hierarchically organized cortico-basal ganglia systems. Hippocampus. 2020 Jan;30(1):73–98. doi: 10.1002/hipo.23167. Epub 2019 Oct 16. PMID: 31617622; PMCID: PMC6972576.

Scharfman HE, Kunkel DD, Schwartzkroin PA. Synaptic connections of dentate granule cells and hilar neurons: results of paired intracellular recordings and intracellular horseradish peroxidase injections. Neuroscience. 1990;37(3):693–707. doi: 10.1016/0306-4522(90)90100-i. PMID: 2247219.

Senzai Y, Buzsáki G. Physiological Properties and Behavioral Correlates of Hippocampal Granule Cells and Mossy Cells. Neuron. 2017 Feb 8;93(3):691–704.e5. doi: 10.1016/j.neuron.2016.12.011. Epub 2017 Jan 26. PMID: 28132824; PMCID: PMC5293146.

Senzai Y. Function of local circuits in the hippocampal dentate gyrus-CA3 system. Neurosci Res. 2019 Mar;140:43–52. doi: 10.1016/j.neures.2018.11.003. Epub 2018 Nov 5. PMID: 30408501.

Shen J, Yao PT, Ge S, Xiong Q. Dentate granule cells encode auditory decisions after reinforcement learning in rats. Sci Rep. 2021 Jul 13;11(1):14360. doi: 10.1038/s41598-021-93721-8. PMID: 34257342; PMCID: PMC8277790.

Song D, Wang D, Yang Q, Yan T, Wang Z, Yan Y, Zhao J, Xie Z, Liu Y, Ke Z, Qazi TJ, Li Y, Wu Y, Shi Q, Lang Y, Zhang H, Huang T, Wang C, Quan Z, Qing H. The lateralization of left hippocampal CA3 during the retrieval of spatial working memory. Nat Commun. 2020 Jun 9;11(1):2901. doi: 10.1038/s41467-020-16698-4. PMID: 32518226; PMCID: PMC7283476.

Sosa M, Giocomo LM. Navigating for reward. Nat Rev Neurosci. 2021 Aug;22(8):472–487. doi: 10.1038/s41583-021-00479-z. Epub 2021 Jul 6. Erratum in: Nat Rev Neurosci. 2021 Sep;22(9):586. PMID: 34230644; PMCID: PMC9575993.

Sutton, R. S., & Barto, A. G. (2018). Reinforcment Learning. Cambridge, MA: The MIT Press.

Svensson, M., Grahm, M., Ekstrand, J., Hoglund, P., Johansson, M., & Tingstrom, A. (2016). Effect of electroconvulsive seizures on cognitive flexibility. Hippocampus, 26(7), 899–910. doi:10.1002/hipo.22573.

L.W. Swanson, et al. An autoradiographic study of the organization of intrahippocampal association pathways in the rat J. Comp. Neurol., 181 (1978), pp. 681–715

Thierry, A. M., Gioanni, Y., Dégénétais, E., and Glowinski, J. (2000). Hippocampo-prefrontal cortex pathway: anatomical and electrophysiological characteristics. Hippocampus 10, 411–419. doi: 10.1002/1098-1063(2000)10:4<411::aid-hipo7>3.0.co;2-a

Tolman, E. C. (1948). Cognitive maps in rats and men. Psychological Review, 55(4), 189–208. 10.1037/h0061626

Treves A, Rolls ET. Computational analysis of the role of the hippocampus in memory. Hippocampus. 1994 Jun;4(3):374–91. doi: 10.1002/hipo.450040319. PMID: 7842058.

Trouche S, Koren V, Doig NM, Ellender TJ, El-Gaby M, Lopes-Dos-Santos V, Reeve HM, Perestenko PV, Garas FN, Magill PJ, Sharott A, Dupret D. A Hippocampus-Accumbens Tripartite Neuronal Motif Guides Appetitive Memory in Space. Cell. 2019 Mar 7;176(6):1393–1406.e16. doi: 10.1016/j.cell.2018.12.037. Epub 2019 Feb 14. PMID: 30773318; PMCID: PMC6424821.

Vaidya AR, Pujara MS, Petrides M, Murray EA, Fellows LK. Lesion Studies in Contemporary Neuroscience. Trends Cogn Sci. 2019 Aug;23(8):653–671. doi: 10.1016/j.tics.2019.05.009. Epub 2019 Jul 3. PMID: 31279672; PMCID: PMC6712987.

van Dijk MT, Fenton AA. On How the Dentate Gyrus Contributes to Memory Discrimination. Neuron. 2018 May 16;98(4):832–845.e5. doi: 10.1016/j.neuron.2018.04.018. Epub 2018 May 3. PMID: 29731252; PMCID: PMC6066591.

Visser E, Matos MR, van der Loo RJ, Marchant NJ, de Vries TJ, Smit AB, van den Oever MC. A persistent alcohol cue memory trace drives relapse to alcohol seeking after prolonged abstinence. Sci Adv. 2020 May 6;6(19):eaax7060. doi: 10.1126/sciadv.aax7060. PMID: 32494694; PMCID: PMC7202866.

Voorn P, Vanderschuren LJ, Groenewegen HJ, Robbins TW, Pennartz CM. Putting a spin on the dorsal-ventral divide of the striatum. Trends Neurosci. 2004 Aug;27(8):468–74. doi: 10.1016/j.tins.2004.06.006. PMID: 15271494.

Pennartz CM, Groenewegen HJ, Lopes da Silva FH. The nucleus accumbens as a complex of functionally distinct neuronal ensembles: an integration of behavioural, electrophysiological and anatomical data. Prog Neurobiol. 1994 Apr;42(6):719–61. doi: 10.1016/0301-0082(94)90025-6. PMID: 7938546.

Weeden CS, Hu NJ, Ho LU, Kesner RP. The role of the ventral dentate gyrus in olfactory pattern separation. Hippocampus. 2014 May;24(5):553–9. doi: 10.1002/hipo.22248. Epub 2014 Jan 28. PMID: 24449260.

Wesnes KA, Annas P, Basun H, Edgar C, Blennow K. Performance on a pattern separation task by Alzheimer’s patients shows possible links between disrupted dentate gyrus activity and apolipoprotein E ∈4 status and cerebrospinal fluid amyloid-β42 levels. Alzheimers Res Ther. 2014 Apr 15;6(2):20. doi: 10.1186/alzrt250. PMID: 24735568; PMCID: PMC4054957.

Witter MP. The perforant path: projections from the entorhinal cortex to the dentate gyrus. Prog Brain Res. 2007;163:43–61. doi: 10.1016/S0079-6123(07)63003-9. PMID: 17765711.

Yassa MA, Stark CE. Pattern separation in the hippocampus. Trends Neurosci. 2011 Oct;34(10):515–25. doi: 10.1016/j.tins.2011.06.006. Epub 2011 Jul 23. PMID: 21788086; PMCID: PMC3183227.

Yun S, Soler I, Tran FH, Haas HA, Shi R, Bancroft GL, Suarez M, de Santis CR, Reynolds RP, Eisch AJ. Behavioral pattern separation and cognitive flexibility are enhanced in a mouse model of increased lateral entorhinal cortex-dentate gyrus circuit activity. Front Behav Neurosci. 2023 Jun 1;17:1151877. doi: 10.3389/fnbeh.2023.1151877. PMID: 37324519; PMCID: PMC10267474.

Zaidel DW. The case for a relationship between human memory, hippocampus and corpus callosum. Biol Res. 1995;28(1):51–7. PMID: 8728820.

Zelikowsky M, Bissiere S, Hast TA, Bennett RZ, Abdipranoto A, Vissel B, Fanselow MS. Prefrontal microcircuit underlies contextual learning after hippocampal loss. Proc Natl Acad Sci U S A. 2013 Jun 11;110(24):9938–43. doi: 10.1073/pnas.1301691110. Epub 2013 May 15. PMID: 23676273; PMCID: PMC3683762.

Zhu H, Yan H, Tang N, Li X, Pang P, Li H, Chen W, Guo Y, Shu S, Cai Y, Pei L, Liu D, Luo MH, Man H, Tian Q, Mu Y, Zhu LQ, Lu Y. Impairments of spatial memory in an Alzheimer’s disease model via degeneration of hippocampal cholinergic synapses. Nat Commun. 2017 Nov 22;8(1):1676. doi: 10.1038/s41467-017-01943-0. PMID: 29162816; PMCID: PMC5698429.

