## Supplementary figures and images for "Transient DREADD manipulation of the dorsal Dentate Gyrus in rats impairs disambiguation of similar place-outcome associations"

### Supplementary Figure 1

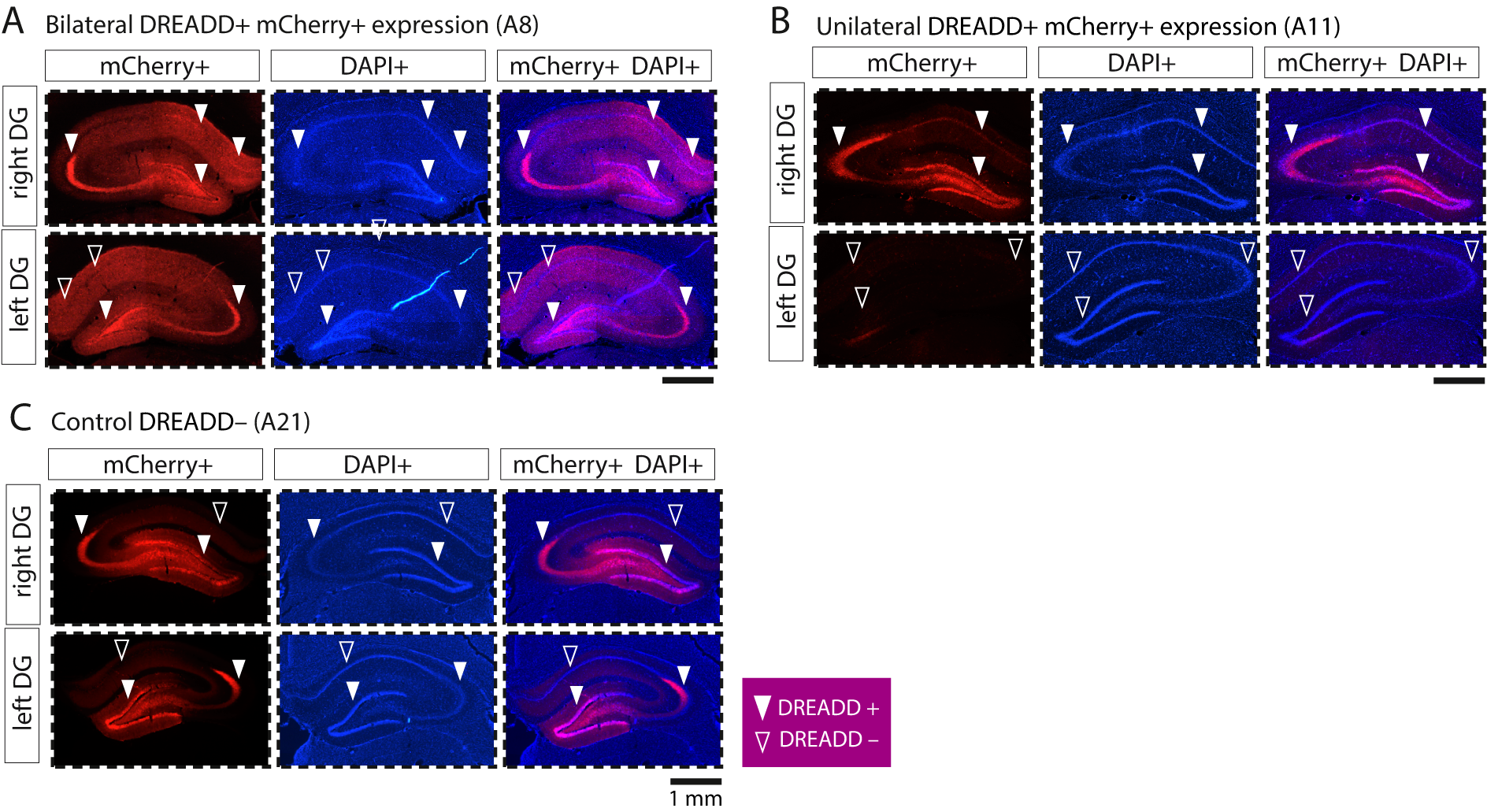

### Supplementary Figure 2

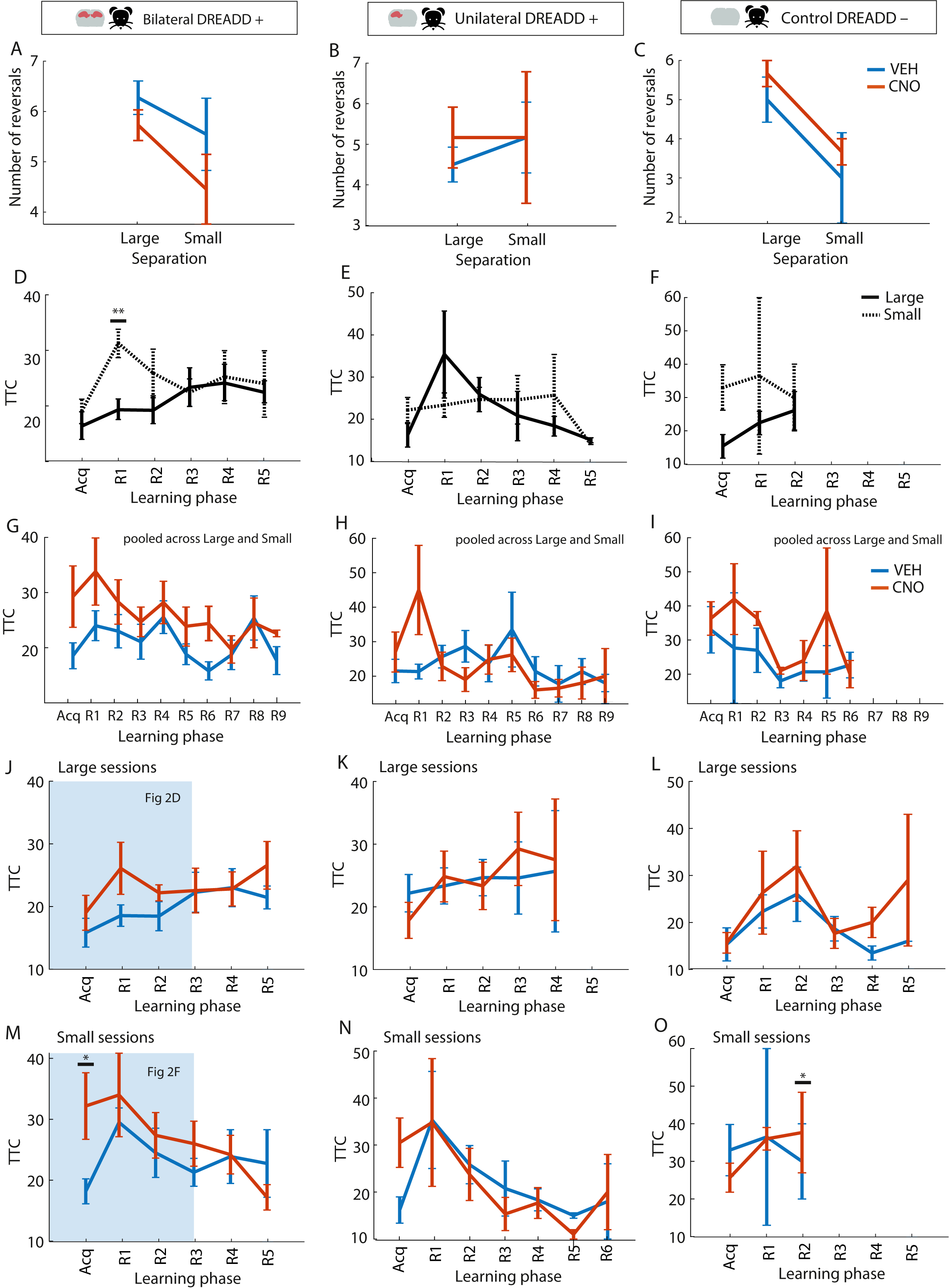

### Supplementary Figure 3

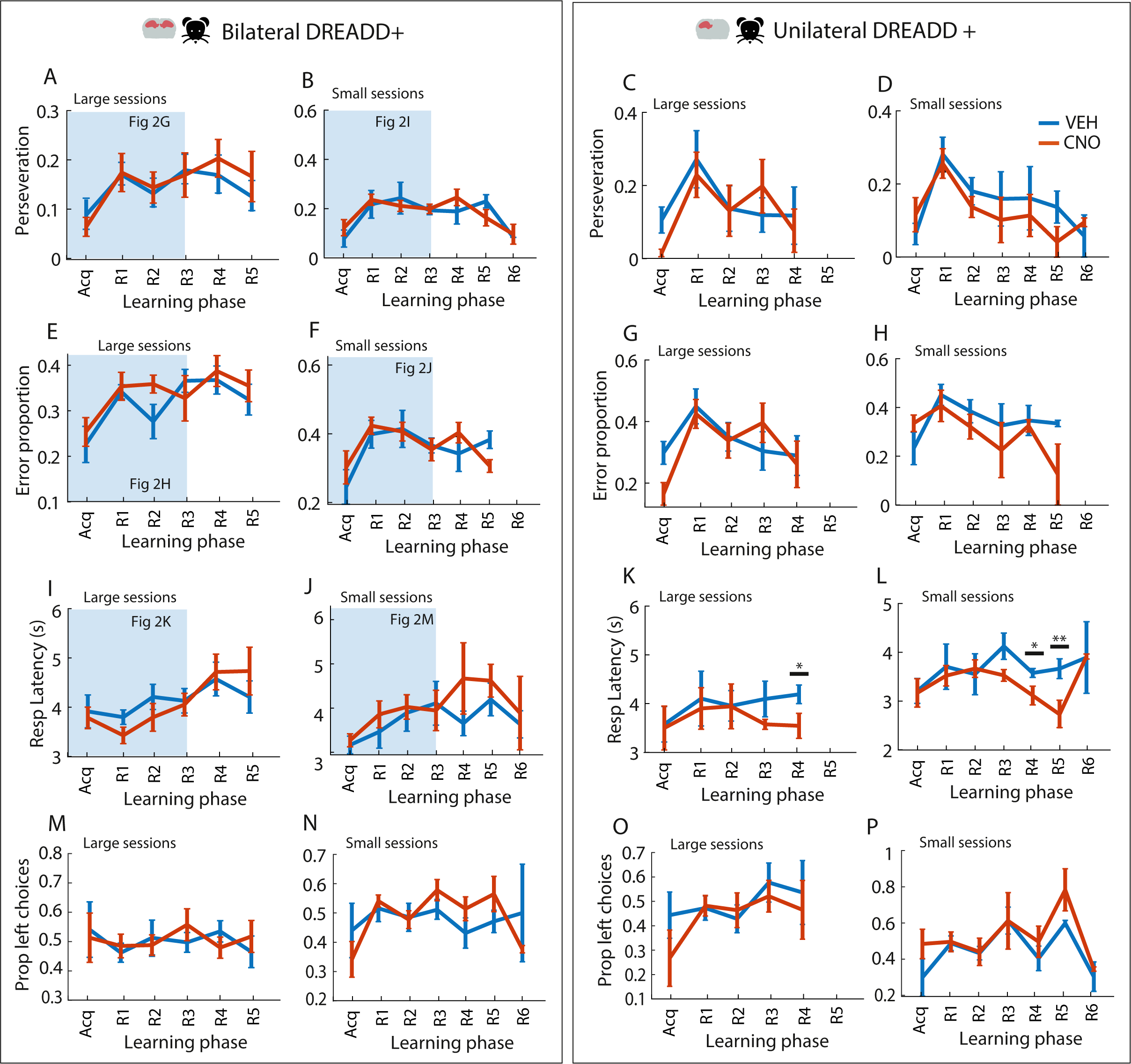

### Supplementary Figure 4

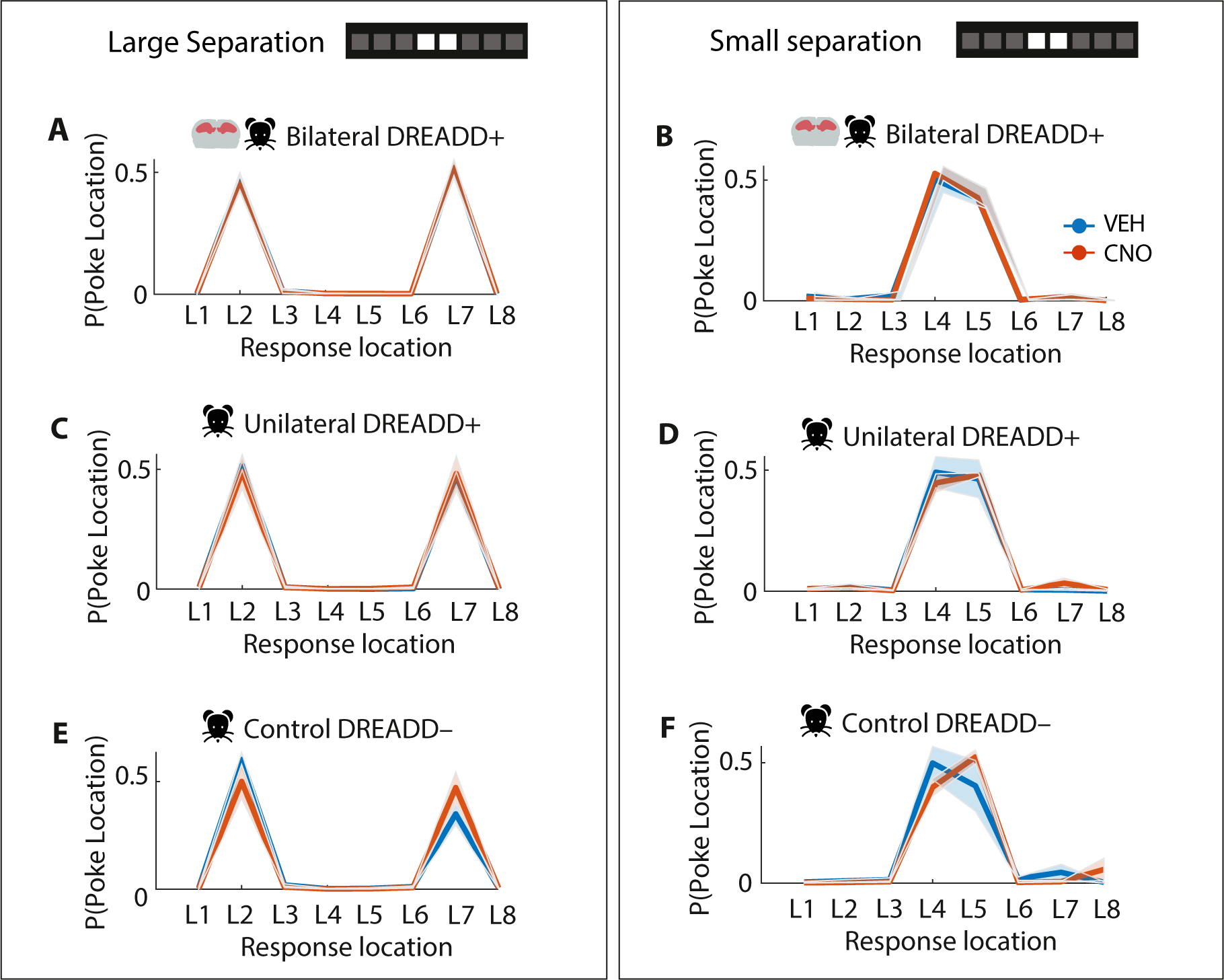

### Supplementary Figure 5

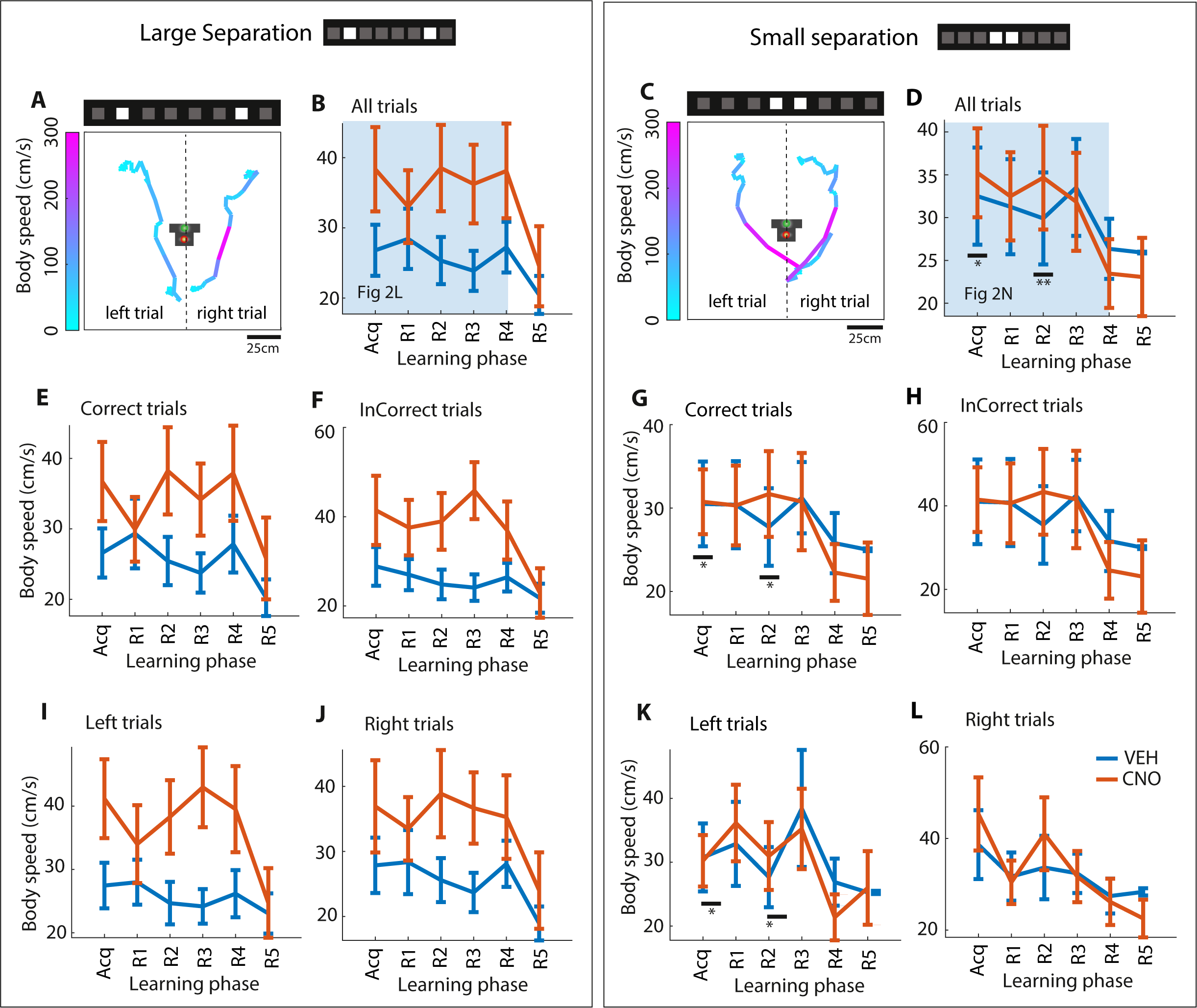

### Supplementary Figure 6

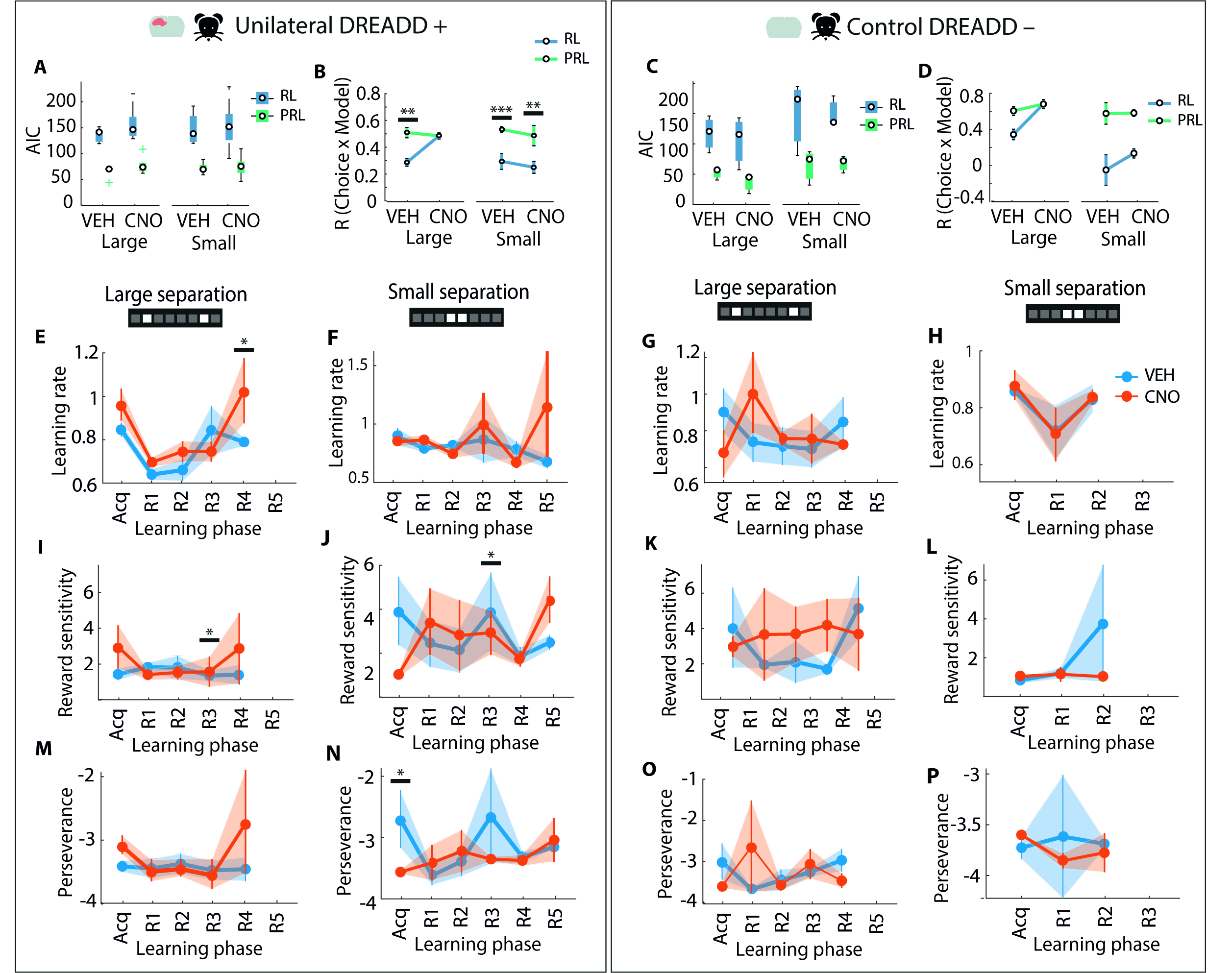
